# Features extracted using tensor decomposition reflect the biological features of the temporal patterns of human blood multimodal metabolome

**DOI:** 10.1101/2022.05.24.493017

**Authors:** Suguru Fujita, Yasuaki Karasawa, Ken-ichi Hironaka, Y-h. Taguchi, Shinya Kuroda

**Author notes:** **Corresponding Author:** Shinya Kuroda, 7-3-1 Hongo, Bunkyo-ku, Tokyo 113, 0033, Japan.

## Abstract

High-throughput omics technologies have enabled the profiling of entire biological systems. For the biological interpretation of such omics data, two analyses, hypothesis- and data-driven analyses including tensor decomposition, have been used. Both analyses have their own advantages and disadvantages and are mutually complementary; however, a direct comparison of these two analyses for omics data is poorly examined.We applied tensor decomposition (TD) to a dataset representing changes in the concentrations of 562 blood molecules at 14 time points in 20 healthy human subjects after ingestion of 75 g oral glucose. We characterized each molecule by individual dependence (constant/variable) and time dependence (sustained/transient). Three of the four features extracted by TD were characterized by our previous hypothesis-driven study, indicating that TD can extract some of the same features obtained by hypothesis-driven analysis in a non-biased manner. In contrast to the years taken for our previous hypothesis-driven analysis, the data-driven analysis in this study took days, indicating that TD can extract biological features in a non-biased manner without the time-consuming process of hypothesis generation.

**Author Summary:** For biological interpretation of lage-scale omics data, two analyses, hypothesis-driven analysis and data-driven analysis including tensor decomposition, have been used. These two analyses have their own advantages and disadvantages, and are mutually complementary. However, the direct comparison between these two analyses for omic data is poorly examined. In this study, we applied tensor decomposition to a dataset representing temporal changes in the human 562 blood molecules as data-driven analysis and extracted three features. We have previously analyzed the same data by hypothesis-driven analysis (Fujita et al., 2022). The three features extracted by the tensor decomposition are the same features extracted by the hypothesis-driven analysis, indicating that the tensor decomposition can extract the features in an unbiased manner. Although the same features can be extracted by the tensor decomposition and hypothesis-driven analysis, hypothesis-driven analysis in our earlier study took years (Fujita et al., 2022), while feature extraction by tensor decomposition took only days in this study. Thus, tensor decomposition can extract biological features in a non-biased manner without time-consuming process of hypothesis generation. We propose that tensor decomposition can be the first choice for analysis of omic data.

## Introduction

The introduction of high-throughput technologies has enabled the profiling of entire biological systems by acquiring omics data such as genomes, transcriptomes, epigenomes, and metabolomes (1,2). The biological interpretation of such omics data requires integrating, summarizing, and visualizing the omics data to acquire a complete picture of the biological system (3). A variety of bioinformatics methodologies have been developed to address the challenge of processing large amounts of complex omics data (4–7). There are two analyses used for omics analyses: hypothesis-driven and data-driven (6,8). Hypothesis-driven analysis tests the hypothesis made by the researcher, whereas data-driven analysis does not require a hypothesis to be made by the researcher in advance. Hypothesis-driven analysis is a subjective analysis that extracts the features of omics data by intuition. This result and its biological interpretation are direct and easy to understand. However, because hypothesis-driven analysis is a human-task, hypothesis generated by the same data may differ between individuals, and the extraction of features can be biased depending on prior knowledge. Also, hypothesis generation relies on human inspections and trial and error, which makes it time-consuming. Data-driven analysis is an objective analysis that extracts features by statistical analysis. Because data-driven analysis is a computational task, feature extraction (FE) can avoid bias from individuals and prior knowledge and is much faster than hypothesis-driven analysis. However, the extracted feature is not necessarily easy to understand and it is sometimes difficult to interpret the biological data. Hypothesis- and data-driven analyses have their own advantages and disadvantages, and are mutually complementary. However, a direct comparison between these two analyses is poorly examined.

We previously used hypothesis-driven analysis for time series data of various blood metabolites such as amino acids and lipids, including blood glucose and hormones during oral glucose ingestion, and found four features with temporal patterns among individuals and molecules (6). However, because the number of molecules targeted was limited and different statistical methods were used to calculate the features, analyst bias is a concern. In addition, the FE took years. To address these issues, we attempted a non-biased FE using tensor decomposition (TD).

Here, we used a data-driven approach based on TD as a multivariate analysis method applied to multi-omics datasets. TD enables data-driven analyses such as data dimensionality reduction, classification, and potential FE (9,10) and has been widely applied to omics studies (Alter and Golub, 2005; Dyrby et al., 2005; Omberg, Golub and Alter, 2007; Yener et al., 2008; Conesa et al., 2010; Acar, Bro and Smilde, 2015; Gardlo et al., 2016; Fanaee-T and Thoresen, 2019; Yahyanejad, 2019

(7,11–16). Omberg *et al.* (2007) integrated genome-scale mRNA expression data from three cell cycle time courses in yeast to identify genes and the differential effects of gene-mediated biological processes on cell cycle progression using TD. TD also has been applied to analyses within each omics (17–21) and among multiple omics (3,22,23). Recently, the application of TD-based unsupervised FE was proposed (Taguchi, 2017a, 2017b, 2017c; Taguchi and Turki, 2019, 2021; Taguchi, 2020). Taken together, the data-driven analyses using TD in these studies have shown that biologically meaningful features can be extracted by a data-driven non-biased method from datasets with various modes. However, TD has not been applied to a time series dataset of human blood metabolites and hormone concentrations before and after glucose ingestion.

In this study, we applied TD to time series datasets representing changes in the concentrations of 562 molecules (555 human blood metabolites and 7 hormones) in 20 healthy subjects before and at 14 time points after ingestion of 75 g oral glucose to extract features. We obtained the core tensor and individual-, time-, and molecule-related singular vectors. Reconstructing the time series using only the dominant singular vectors allowed us to better interpret the features of the temporal patterns of the molecules. We characterized each molecule by individual dependence (constant or variable) and time dependence (sustained or transient). The molecule-related singular vectors obtained by TD reflected three of the four features characterized by our previous hypothesis-driven study (6). We also extracted 68 molecules showing feature time and individual dependencies through the unsupervised learning method by using TD. The extracted molecules significantly overlapped with the analyzed molecules in our previous study (6). Therefore, by applying TD to the dataset characterized in our earlier study, we were able to extract the features of the target molecules and reveal the temporal patterns in that study (6). This result not only confirms the validity of our previous findings but also shows the usefulness of TD as a FE method.

Next, we applied the TD method to a dataset representing the concentration changes of 40 molecules in three healthy subjects at 26 time points by three different oral doses (25g, 50g, 75g) and two different patterns of glucose ingestion (bolus or 2h-continuous ingestion). We obtained the core tensor and individual-, time-, molecule-, and experimental condition-related singular vectors. Reconstructing the time series using only the dominant singular vectors allowed us to better interpret the features of the temporal patterns of the molecules. We characterized the temporal pattern of each molecule by its experimental condition dependence (constant or variable) and time dependence (early or late peaks). Thus, we applied TD to a time series dataset of human blood metabolites and hormone concentrations before and after glucose ingestion with various modes, and extracted the features in a non-biased manner. Of the four features of temporal patterns in the hypothesis-driven analysis from our previous study (6), the following three features were extracted by data-driven analysis: the “amplitude and rate” components of temporal patterns, the similarity of temporal patterns among individuals, and the similarity of temporal patterns among molecules. The only feature not extracted was the relationship among individuals over time. This resultindicates that TD can extract some of the same features obtained by hypothesis-driven analysis in a non-biased manner. In addition, FE using TD took only days in this study. As we extracted biological features in a non-biased manner without the time-consuming process of hypothesis generation, we propose that TD be the first choice for the analysis of omics data.

## Methods

### Subjects

The study included 20 healthy subjects for ‘*third-order tensor with individual mode, time mode, and molecule mode*’ and 3 healthy subjects for ‘*fourth-order tensor with individual mode, time mode, experimental condition mode, and molecule mode.*’ The subjects’ profiles are shown in Table S1, and all subjects provided informed consent. Subjects were recruited through a snowball sampling and self-selection.

### Blood sampling experiment

We used human blood samples obtained previously (Fujii et al., 2019, 2022). Briefly, after a 10 h overnight fast, subjects underwent the oral glucose tolerance test in the morning. An intravenous catheter was inserted into the vein of the forearm and fasting samples were drawn twice.

#### ‘third-order tensor with individual mode, time mode, and molecule mode’

A glucose solution containing 75 g glucose (TRELAN-G75^®^; Ajinomoto Inc., Tokyo, Japan) or the same amount of water was orally ingested within a few minutes. Blood samples were obtained at 10, 20, 30, 45, 60, 75, 90, 120, 150, 180, 210, 240 min after ingestion as previously described (Fujita et al., 2022).

#### ‘fourth-order tensor with individual mode, time mode, experimental condition mode, and molecule mode’

A glucose solution containing 25, 50, or 75 g glucose was orally ingested. The ingestion method was rapid within a minute (bolus ingestion) and continuous over the course of 2 h (2 h continuous ingestion). For continuous ingestion, we connected the tube to a noncontact microdispenser robot (Mr. MJ; MECT Co., Osaka, Japan) (25). The glucose solution was ingested from the tube, and blood samples were obtained every 10 min until 240 min after sugar ingestion (26). Subjects remained at rest throughout the test. Blood samples were rapidly centrifuged.

### Sample preparation and measurement

Sample preparation and measurement were performed as previously described (Fujii et al., 2019, 2022). Plasma (40 μL) was extracted with the addition of 400 μL ice-cold methanol containing internal standards (10 mM L-methionine sulfone [Wako, Tokyo, Japan], 100 mM 2-morpholinoethanesulfonic acid [Dojindo Molecular Technologies, Rockville, MD, USA], 100 mM D-10-camphorsulfonic acid [Wako]), 400 μL chloroform, and 120 μL water. After centrifugation at 10,000 × *g* for 3 min at 4°C, the separated aqueous layer was filtered through a 5 kDa cutoff filter (Millipore, Burlington, MA, USA) to remove protein contamination. The filtrate (300 μL) was lyophilized and dissolved in 20 μL water containing two reference compounds (200 μM each of trimesate [Wako] and 3-aminopyrrolidine [Sigma-Aldrich, St. Louis, MO, USA]) for migration time and then injected into a capillary electrophoresis time-of-flight mass spectrometry system (Agilent Technologies, Santa Clara, CA, USA) (27–29). Among the measured molecules, gastric inhibitory polypeptide (GIP) (active) was measured using an enzyme-linked immunosorbent assay kit. Blood hormones and some metabolites were measured according to methods developed by LSI Medience Co., Ltd. (Tokyo, Japan).

### Ethics committee certification

We complied with Japan’s Ethical Guidelines for Epidemiological Research, and the study was approved by the institutional review board and ethics committee of Tokyo University Hospital (No. 10264-(4)).

### Data preprocessing

Because we focused on temporal patterns in this study for ‘*third-order tensor with individual mode, time mode, and molecule mode,’* we included only metabolites and hormones that we measured to obtain our time series data. Thus, we excluded the molecules included in Table S2 from our TD analysis. We considered the missing values to be zero. For ‘*third-order tensor with individual mode, time mode, and molecule mode,’* we considered the mean value to be −10 and the fasting value to be 0 min; and for ‘*fourth-order tensor with individual mode, time mode, experimental condition mode, and molecule mode,’* we considered the mean value to be −5 and the fasting value to be 0 min.

### TD-based unsupervised FE

For ‘*third-order tensor with individual mode, time mode, and molecule mode*,’ we chose *w*_*l*_3_*k*_ to select biologically meaningful molecules. After selecting *w*_*l*_3_k_, assuming that *w*_*l*_3_k_ were normally distributed, the *p*-values were attributed to the *k* th molecules using the *χ* squared distribution as:

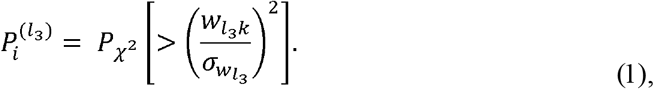

where *σ*_*w*_*l*_3___ is the standard deviation of *w*_*l*_3_*k*_, and *P* χ ^2^[> *x*] is the probability that the argument is larger than *x* under the assumption that the arguments obey a *χ* squared distribution. The *p*-values were further adjusted by the Benjamini-Hochberg (BH) criterion (24), and those molecules associated with adjusted *p* < 0.1 were selected.

### Reconstructed time courses

For *‘third-order tensor with individual mode, time mode, and molecule mode,’* the original time series of the molecule was reconstructed using only constant individual dependence (*u*_*l*_1_*i*_, *l*_1_ = 1) and variable individual dependence (*u*_*l*_1_*i*_, *l*_1_ = 1) (Fig. 3). For “individuals similar pattern” and “individuals different pattern,” the concentration of the *i* th individual of the *j* th time point of the *k*th molecule is defined as follows:

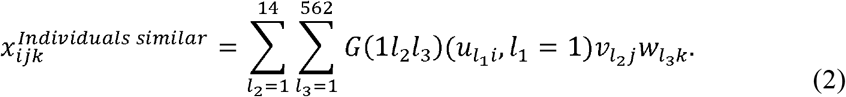

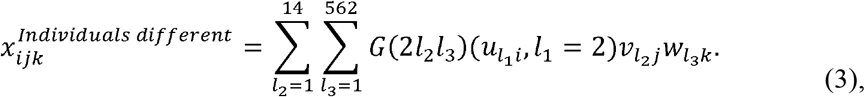

where (1 *or* 2*l*_2_*l*_3_) ∈ ℝ^20×14×562^ is the core tensor, *U* = {*u*_l_1_*i*_} ∈ ℝ^20×20^, *V* = {*v*_*l*_2_*j*_} ∈ ℝ^14×14^,and *W* = {*w*_*l*_3_*k*_} ∈ ℝ^562×562^ are the orthogonal matrices.

For *‘fourth-order tensor with individual mode, time mode, experimental condition mode, and molecule mode,’* the original time series of the molecule was reconstructed using only constant experimental condition dependence (*w*_*l*_3_*k*_, *l*_3_ = 1) and variable individual dependence (*w*_*l*_3_*k*_, *l*_3_ = 1) (Fig. 7). For “conditions similar pattern” and “conditions different pattern,” the concentration of the *i*th individual’s of the *j*th time point of the *k*th molecule of the *m*th experimental condition is defined as follows:

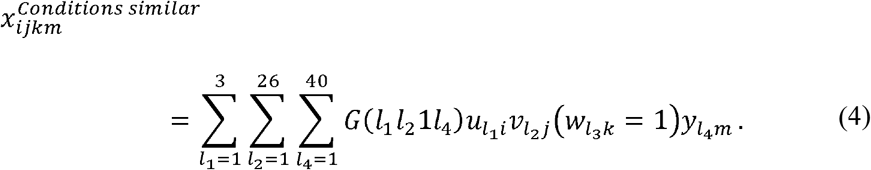

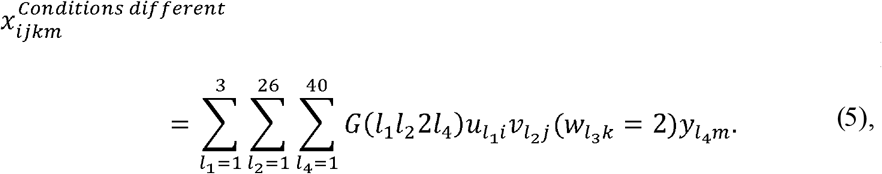

where *G*(*l*_1_*l*_2_1 *or* 2*l*_4_) ∈ ℝ^3×26×6×40^ is the core tensor, *U* = {*u*_*l*_1_*i*_} ∈ ℝ^3×3^, *V* = {*v*_*l*_2_*j*_} ∈ ℝ^26×26^, *W* = {*w*_*l*_3_*k*_} ∈ ℝ^6×6^, *Y* = {*y*_*l*_4_*m*_} ∈ ℝ^10×10^ are the orthogonal matrices.

## Results

### Introduction of TD (*‘third-order tensor with individuals mode, time mode, and molecule mode*’)

We applied TD to the multimodal data of 562 blood molecules after ingestion of 75 g oral glucose at 14 time points in 20 healthy subjects (Fig. 1). Multimodal data are a data structure X with three axes (modes): molecules, individuals, and time. We applied a TD called Tucker decomposition. We schematically illustrate the Tucker decomposition model for the three-dimensional data case in Figure 1. The data cube X (*X* ∈ ℝ^20×14×562^) was decomposed into several components by Tucker decomposition, and the number of components differed in the three modes (i.e., dimensions or axes). Here, *G*(*l*_1_*l*_2_*l*_3_ ∈ ℝ^20×14×562^ is the core tensor and *U* ∈ ℝ^562×562^, *V* ∈ ℝ^20×20^, *W* ∈ ℝ^14×14^ are the orthogonal matrices. The extracted components were characterized by the columns of the three orthogonal matrices (*U, V, W*) (i.e., the singular vectors of each mode), meaning that the extracted components had a dependence on each column of the original data. The model of the original data is a weighted sum of the outer products among the columns of (*U, V, W*). The core tensor G represents the weighted value of the product of single components.

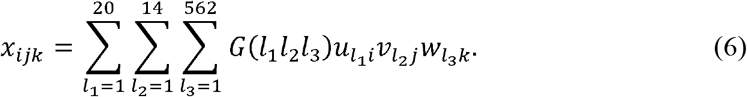

We applied one of the algorithms of the Tucker decomposition, called higher order singular value decomposition (HOSVD). This method is easier to compute and has a more converging algorithm than the other method because it does not need initialization and the iterative computation (24). The components of X (*x_ijk_*) can be decomposed as in (Eq. 1), where *G*(*l*_1_*l*_2_*l*_3_ ∈ ℝ^20×14×562^ is the core tensor; *U* = {*u*_*l*_1_*i*_} ∈ ℝ^20×20^, *V* = {*v*_*l*_2_*j*_} ∈ ℝ^14×14^, *W* = {*w*_*l*_3_*k*_} ∈ ℝ^562×562^ are the orthogonal matrices; *i* means individual; *j* means time point; and *k* means molecule.

**Fig. 1.**
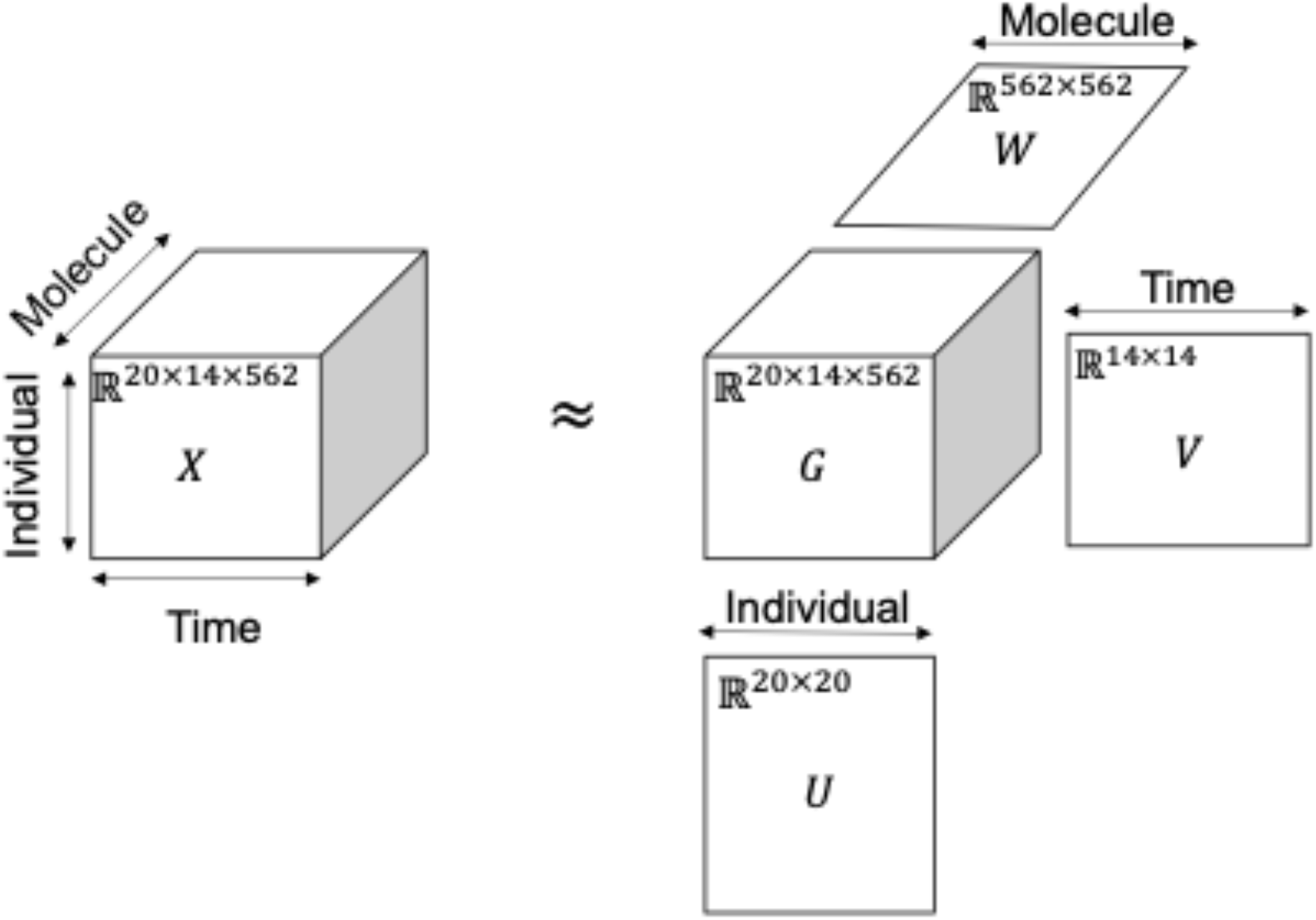
Tensor decomposition using the Tucker model is a weighted sum of outer products between the components stored as columns in *U, V, W*. Twenty healthy subjects orally ingested glucose with 75 g bolus ingestion. The data structure *X* ∈ ℝ^20×14×562^) had three axes: 20 individuals × 14 time points × 562 molecules. *G*(*l*_1_*l*_2_*l*_3_) ∈ ℝ^20×14×562^ is the core tensor and *U* ∈ ℝ^20×20^, *V* ∈ ℝ^14×14^, *W* ∈ ℝ^562×562^ are the singular matrices of individuals, time, and molecules.

### Singular vectors obtained by TD (‘*third-order tensor with individuals mode, time mode, and molecule mode*’)

We performed TD of three modes data of the concentrations of 562 blood molecules (7 hormones, 555 metabolites) at 14 time points in 20 healthy subjects after ingestion of 75 g oral glucose, and attempted to capture the differences in temporal patterns of molecular concentration among individuals and molecules by glucose ingestion with TD. The data are formatted as tensor *X* = }*x_ijk_* ∈ ℝ^20×14×562^ representing the concentration of the *i*th individual of the *j*th time point of the *k*th molecule (Fig. 1). We normalized *x_ijk_* as 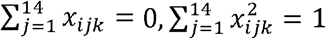. Higher order singular value decomposition (24) (HOSVD) was applied to *x_ijk_*.

To select a set of molecules with specific time and individual dependence, we used the method proposed by Taguchi (2020). In this method, we focused on the absolute value of *G*(*l*_1_*l*_2_*l*_3_) and investigated the combinations of *u*_*l*_1_*i*_, *v*_*l*_2_*j*_, *w*_*l*_3_*k*_ that shared a large absolute value of *G*(*l*_1_*l*_2_*l*_3_). *u*_*l*_1_*i*_, *v*_*l*_2_*j*_, *w*_*l*_3_*k*_ are singular vectors related to individuals, time, and molecules, respectively. This is because we could consider a molecule with a large value of *u*_*l*_1_*i*_, which shares a large absolute value of *G*(*l*_1_*l*_2_*l*_3_) with a specific pair of individual and time singular vectors, as shown in Figure S1B, to have specific individual and time dependencies. Because *u*_*l*_1_*i*_, *v*_*l*_2_*j*_, *w*_*l*_3_*k*_ are the unit vectors, note that *G*(*l*_1_*l*_2_*l*_3_) represents the weighted value of the product of any single component, as well as how much each dependence is related to each other (Eq. 1).

We sorted core tensor components *G*(*l*_1_*l*_2_*l*_3_) in the order of their absolute values (Table 1), focusing only on the top four core tensor components and associated singular vectors (Fig. S1C, Table 1). As discussed later, these four were sufficient for replicating the features of temporal patterns that characterized differences among molecules and individuals in our previous study (6). These large absolute values of *G*(*l*_1_*l*_2_*l*_3_) corresponded to the singular vectors *u*_*l*_1_*i*_, *l*_1_ = 1,2, *v*_*l*_2_*j*_, *l*_2_ = 1,3, *w*_*l*_3_*k*_, *l*_3_ = 1,2,4 of individual, time, and molecule (Table 1). Thus, we next interpreted these singular vectors.

*u*_*l*_1_*i*_ are the individual-related singular vectors; these values mean individual dependence (Fig. 2A, B). *u*_*l*_1_*i*_, *l*_1_ = 1 showed a constant value among individuals (Fig. 2A). For *u*_*l*_1_*i*_, *l*_1_ = 2, the values showed variable among individuals. (Fig. 2B). This result means that *u*_*l*_1_*i*_, *l*_1_ = 1 represents the common component among individuals and *u*_*l*_1_*i*_, *l*_1_ = 2 represents the variable component among individuals, and that the common component among individuals is larger than the variable component among individuals.

**Fig. 2.**
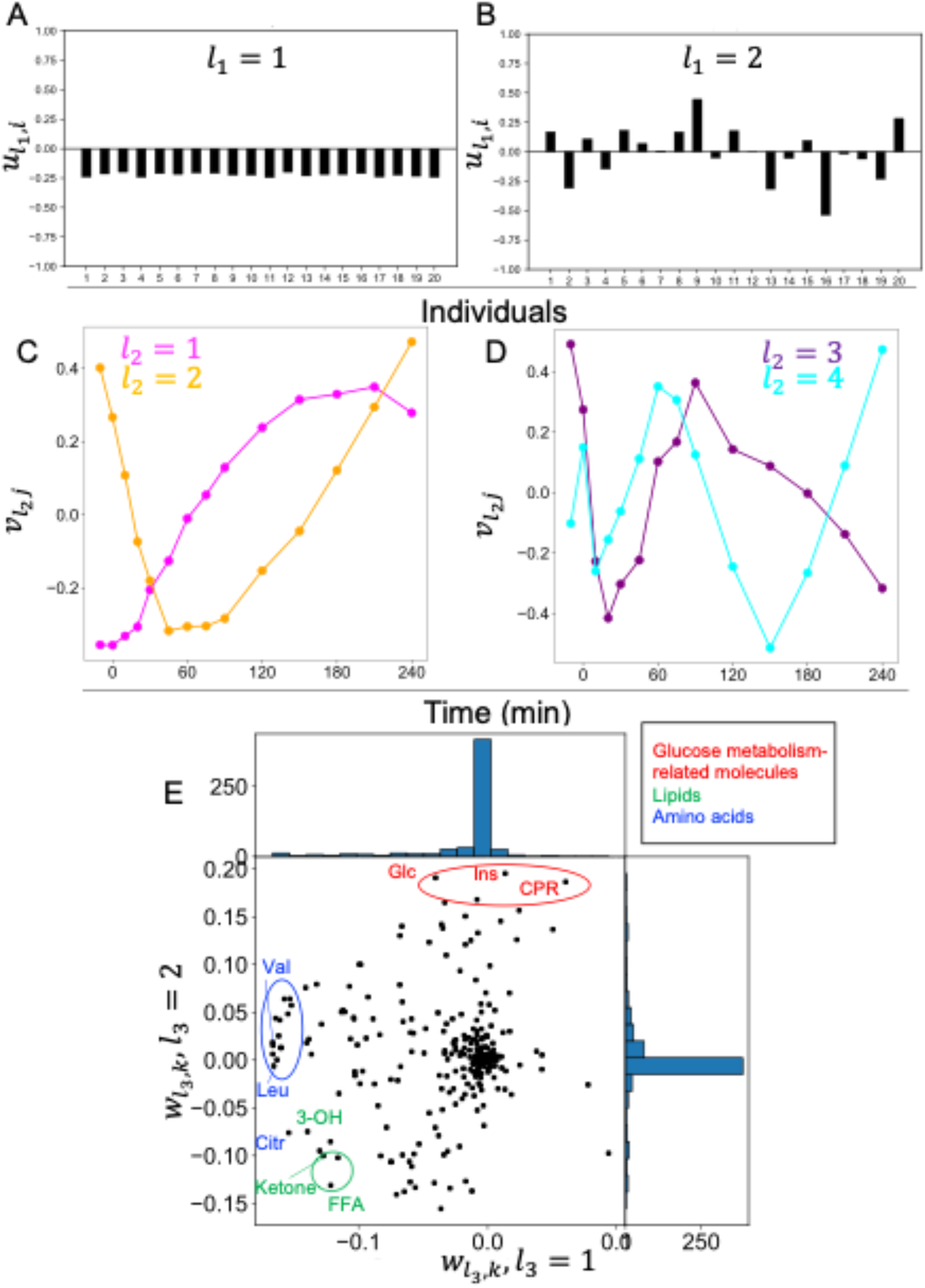
Singular vectors obtained by tensor decomposition (’*third-order tensor with individuals mode, time mode, and molecule mode*’) **A** Individual-related singular vectors (*u*_*l*_1_*i*_, *l*_1_ = 1). **B** Individual-related singular vectors (*u*_*l*_1_*i*_, *l*_1_ = 2). **C** Time-related singular vectors (Magenta line: *v*_*l*_2_*j*_, *l*_2_ = 1 Orange line: *v*_*l*_2_*j*_, *l*_2_ = 2). **D** Time-related singular vectors (Purple line: *v*_*l*_2_*j*_, *l*_2_ = 3, Cyan line: *v*_*l*_2_*j*_, *l*_2_ = 4). **E** Molecule-related singular vectors (*w*_*l*_3_*k*_, *l*_3_ = 1,2). Representative molecules are labeled. Abbreviations for the representative molecules are as follows: Cit, citrulline; CRP, C-reactive peptide; FFA, free fatty acid; 3-OH, 3-hydroxybutyric acid; Ketone, Total ketone body; Glc, glucose; Ins, insulin; Leu, leucine; Val, valine. The label colors correspond to the metabolic group list.

**Fig. 3.**
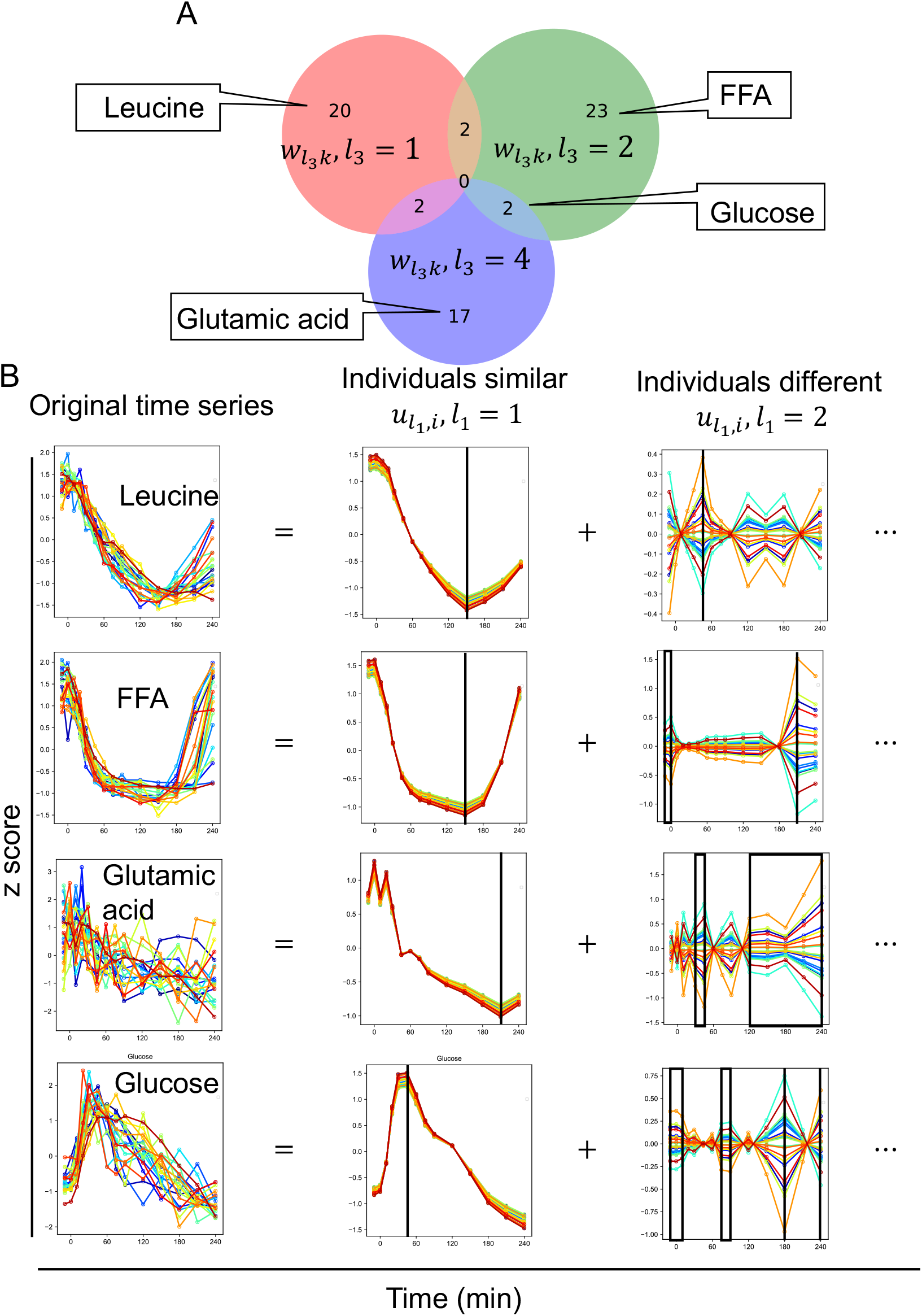
Set of molecules selected by tensor decomposition and decomposed time series by the dominant singular vectors (‘*third-order tensor with individuals mode, time mode, and molecule mode*’) **A** Venn diagram of the set of molecules selected by tensor decomposition. These sets include the molecules whose time series are shown in B. **B** Decomposed time series by the dominant singular vectors. Decomposed time series by other than the dominant singular vectors (***u***_***l***_1_*i*_, *l*_1_ > **2**) are omitted. The black lines in “individuals similar” indicate the time point of the peak. The black boxes and black lines in “individuals different” indicate time points of high variability among individuals. The color of each line indicates each individual. Abbreviations for the molecule are as follows: FFA, free fatty acid.

*v*_*l*_2_*j*_ are the time-related singular vectors (Fig. 2C, D). These values are time-dependent. *v*_*l*_2_*j*_, *l*_2_ = 1 showed a sustained temporal pattern (Fig. 2C), *v*_*l*_2_*j*_, *l*_2_ = 2 showed a transient temporal pattern (Fig. 2C), and *v*_*l*_2_*j*_, *l*_2_ ≥ 3 showed a zigzag temporal pattern with different periods (Fig. 2D). This result indicates that sustained and transient temporal patterns are the main components of temporal changes. We defined transient as the temporal pattern that returned to 75% of the peak in the time series within 240 min after glucose ingestion, and sustained as the other temporal patterns (Fig. S1A).

*w*_*l*_3_*k*_ are the molecule-related singular vectors (Fig. 2E). These values mean molecule dependence. *w*_*l*_3_*k*_, *l*_3_ = 1 showed large values for amino acids such as leucine and valine (Fig. 2E). *w*_*l*_3_*k*_, *l*_3_ = 2 showed large values for glucose metabolism-related molecules such as glucose and insulin (Fig. 2E). Both *w*_*l*_3_*k*_, *l*_3_ = 1,2 showed large values for lipids such as free fatty acid (FFA) and ketone (Fig. 2E). *w*_*l*_3_*k*_, *l*_3_ = 4 showed large values for the amino acid glutamic acid and succinate, which is a component of the tricarboxylic acid cycle (TCA) cycle (Fig. S2A). This result indicates that molecules with similar value share similar individual and time dependencies, whereas molecules with different value have different individual and time dependencies.

We focused on the top four core tensor components *G*(*l*_1_*l*_2_*l*_3_) in terms of absolute values (Fig S1C, Table 1). Following this, *p*-values were attributed to the *i*th molecule for each of *w*_*l*_3_*k*_, *l*_3_ = 1,2,4 (Methods). *P*-values were corrected using the BH criterion (24), resulting in the selection of 20 molecules for *w*_*l*_3_*k*_, *l*_3_ = 1, 23 molecules for *w*_*l*_3_*k*_, *l*_3_ = 2, and 19 molecules for *w*_*l*_3_*k*_, *l*_3_ = 4 (Table S3).

### Set of molecules selected by TD and decomposed time series by the dominant singular vectors (*‘third-order tensor with individuals mode, time mode, and molecule mode*’)

We investigated the overlap among the selected sets of molecules using *w*_*l*_3_*k*_, *l*_3_ = 1, 2, 4, and found that the overlap was small and the time and individual dependencies of each selected set of molecules were specific (Fig. 3A). To better interpret the features of the temporal patterns of the molecules, we reconstructed the time series using only the singular vectors that we focused on (Fig. 3B, see Methods). This is equivalent to a low-rank approximation of the tensor.

Because we were interested in features of the temporal patterns that are similar among individuals of each molecule and the temporal patterns that are different among individuals, we reconstructed the original time series of the molecule using only the constant individual dependence (*u*_*l*_1_*i*_, *l*_1_ = 1) but also the variable individual dependence *u*_*l*_1_*i*_, *l*_1_ = 2) (Fig. 3B, see Methods). Here, we refer to the time series reconstructed using only (*u*_*l*_1_*i*_, *l*_1_, = 1) as “individuals similar pattern,” and the time series reconstructed using only (*u*_*l*_1_*i*_, *l*_1_ = 2) as “individuals different pattern.” We focused on leucine, FFA, glutamic acid, and glucose, which showed significant responses before and after glucose ingestion in our previous study (6).

For leucine, as one of the representative molecules of the amino acid (Fig. 3B), “individuals similar pattern” peaked at 150 min and showed a sustained temporal pattern. “individuals different pattern” showed the most variation at 45 min, but the absolute value of “individuals different pattern” was smaller than “individuals different pattern” for the other molecules (Fig. 3B). Twenty-three molecules selected only by *w*_*l*_3_*k*_, *l*_3_ = 1 included branched chain amino acids such as leucine and valine, and aromatic amino acids such as tyrosine and phenylalanine (Fig. 3, Table 1). These molecules had a constant individual dependence (*u*_*l*_1_*i*_, *l*_1_ = 1) with a sustained time dependence (*v*_*l*_2_*j*_, *l*_2_ = 1) (Fig. 2, Table 1). This result suggests that the amino acids selected only *w*_*l*_3_*k*_, *l*_3_ = 1 had similar sustained temporal patterns among individuals.

For FFA selected only by *w*_*l*_3_*k*_, *l*_3_ = 2, the “individuals similar pattern” showed a transient temporal pattern with a peak at 150 min (Fig. 3), which is similar to leucine. However, the “individuals difference pattern” of FFA showed the most variation at 210 min, and also showed variation between −10 min and 10 min, which differed from leucine. Twenty-five molecules selected only by *w*_*l*_3_*k*_, *l*_3_ = 2 included glucose metabolism-related molecules and lipids such as FFA and acetoacetic acid (Fig. 3, Table S1). These molecules had variable individual dependence (*u*_*l*_2_*j*_, *l*_1_ = 2) with sustained time dependence (*v*_*l*_2_*j*_, *l*_2_ = 1), while constant individual dependence (*u*_*l*_1_*i*_, *l*_1_ = 1) with transient time dependence (*v*_*l*_2_*j*_, *l*_2_ = 2) (Fig. 2, Table 1). This result suggests that the glucose metabolism-related molecules and lipids, which were selected only by *w*_*l*_3_*k*_, *l*_3_ = 2, had similar transient temporal patterns and different sustained temporal patterns among individuals.

For glutamic acid selected only by *w*_*l*_3_*k*_, *l*_3_ = 4, the “individual similar pattern” peaked at 210 min and showed a sustained temporal pattern, which is slower than leucine and FFA. “individual different pattern” of glutamic acid showed a large variation from 120 to 240 min (Fig. 3), and also showed a large variation from 30 to 45 min (Fig. 3), which is different from leucine and FFA. Seventeen molecules selected only by *w*_*l*_3_*k*_, *l*_3_ = 4 included molecules such as citrate, succinate, and malate, which constitute the TCA cycle, and amino acids such as glutamic acid and glutamine (Fig. 3, Table S1). These molecules had a transient time dependence (*v*_*l*_2_*j*_, *l*_2_ = 2) with variable individual dependence (*u*_*l*_1_*i*_, *l*_1_ = 2) (Fig. 2, Table 1). This result suggests that the transient temporal patterns of these molecules are different among individuals. For glucose both selected by both *w*_*l*_3_*k*_, *l*_3_ = 2, “individuals similar pattern” peaked at 50 min and showed a transient temporal pattern, which is faster than leucine, FFA, and glutamic acid. The “individuals difference pattern” of glucose showed the largest variation at 180 min (Fig. 3), and also showed a large variation from −10 to 10 min, from 75 to 90 min, and at 240 min (Fig. 3), which is different from leucine, FFA, and glutamic acid. For the overlap, the molecules selected by both *w*_*l*_3_*k*_, *l*_3_ = 2 and *l*_3_ = 4 were glucose, creatine, and creatinine (Fig. 3, Table S1). This result suggests that these molecules are characterized by both similar transient temporal patterns among individuals and different sustained temporal patterns among individuals; and different transient temporal patterns among individuals.

For “individuals similar patterns,” leucine and glutamic acid showed sustained patterns, whereas FFA, and glucose showed transient patterns (Fig. 3, Table 1). Furthermore, the different temporal patterns among individuals were sustained for glucose-related molecules and lipids (Table 1). For the “individuals different pattern,” the four molecules did not show a clear time dependence, but the time points with large variation were different among molecules (Fig. 3). Taken together, each metabolic group was characterized by individual dependence (constant or variable) and time dependence (sustained or transient).

### Comparison of the results of our previous study with this study

In our previous study, we used hypothesis-driven analysis and characterized the temporal patterns among individuals and molecules by hypothesis-driven analysis with the following four features (6): the decomposability into “amplitude” and “rate” components, the similarity of temporal patterns among individuals, the relationship among individuals’ over time, and the similarity of temporal patterns among molecules (Fig. S3). In this study, we used TD as data-driven analysis. We investigated whether the features extracted by data-driven analysis in this study reflected the four features derived from the hypothesis-driven analysis. Here, we used the value of the square of *w*_*l*_3_*k*_ to select the molecules.

In our earlier study of the hypothesis-driven analysis, we decomposed the temporal pattern of the molecule into “amplitude of response (AUC)” and “rate of response (T_AUC1/2_)” as the first feature (6). The larger the value of AUC positively or negatively, the larger the response from fasting after glucose ingestion positively (increase) or negatively (decrease), respectively. The larger the value of T_AUC1/2_, the slower the response. In this data-driven analysis, we used *w*_*l*_3_*k*_, *l*_3_ = 1,2,4 to select molecules whose time dependence was transient or persistent. In other words, *w*_*l*_3_*k*_, *l*_3_ = 1,2,4 are likely to capture the features of the temporal pattern such as the amplitude and rate of response. Therefore, we investigated the relationship between the values of *w*_*l*_3_*k*_, *l*_3_ = 1,2,4 and AUC, T_AUC1/2._ Correlation analysis showed a significant relationship between the value of abs (*w*_*l*_3_*k*_, *l*_3_ = 1) and the amplitude of the response, and between the value of abs (*w*_*l*_3_*k*_, *l*_3_ = 2) and the rate of the response, respectively (Fig. 4A, B, Table 2). Thus, the value of *w*_*l*_3_*k*_, *l*_3_ = 1 was negatively correlated with the amplitude of response, and the value of *w*_*l*_3_*k*_, *l*_3_ = 2 was negatively correlated with the rate of response, indicating that these two values extracted by data-driven analysis reflected the features of amplitude and rate of response derived from the hypothesis-driven analysis.

**Fig. 4.**
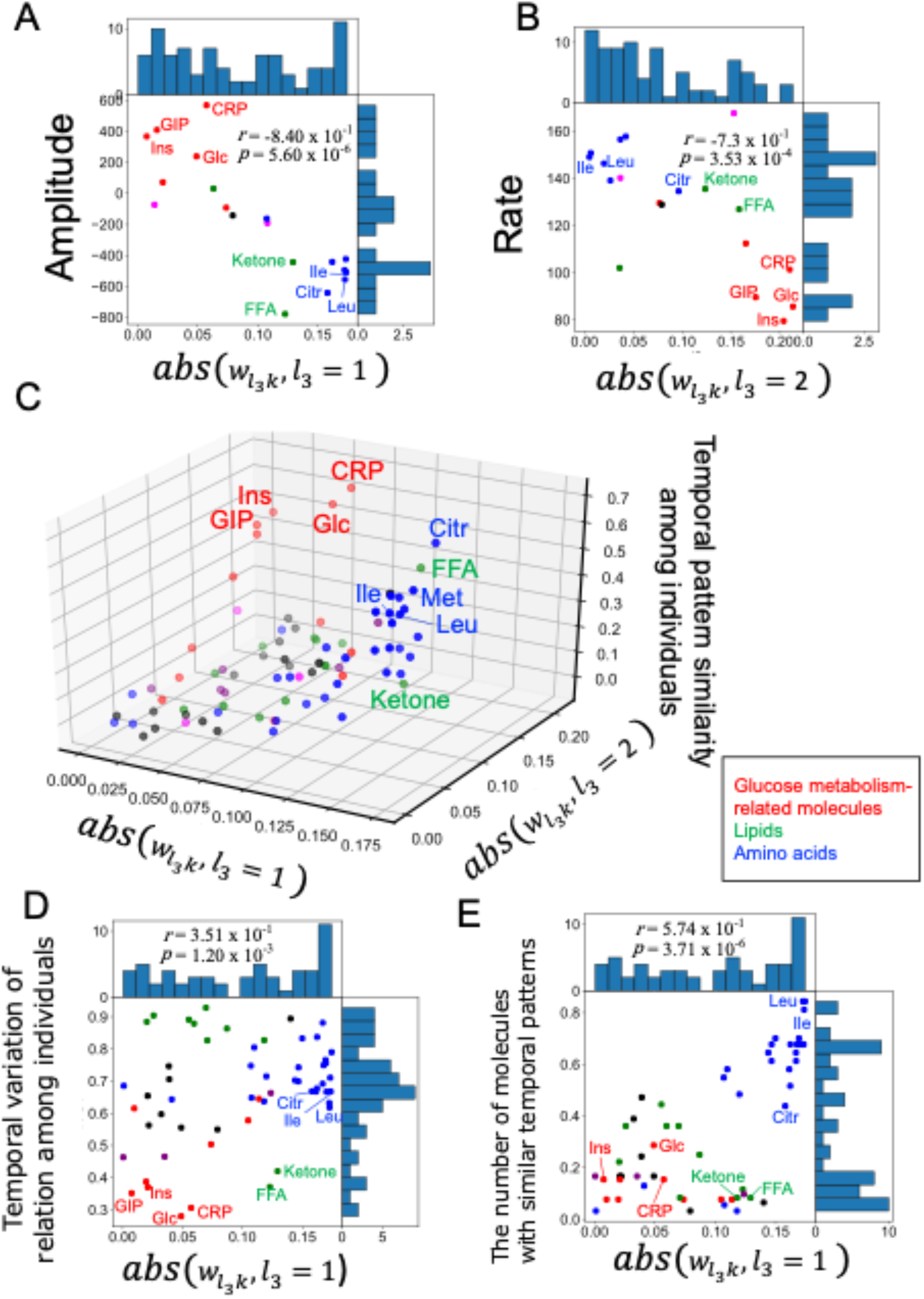
Comparison of tensor decomposition analysis with hypothesis-driven analysis. **A** The distribution of abs (*w*_*l*_3_*k*_, *l*_3_ = 1) and magnitude of response for each molecule. **B** The distribution of abs (*w*_*l*_3_*k*_, *l*_3_ = 2) and rate of response for each molecule. **C** The distribution of abs (*w*_*l*_3_*k*_, *l*_3_ = 1), abs (*w*_*l*_3_*k*_, *l*_3_ = 2) and the similarity of temporal pattern among individuals for each molecule. **D** The distribution of abs (*w*_*l*_3_*k*_, *l*_3_ = 1) and the change in the relationship among individuals over time for each molecule. **E** The distribution of abs (*w*_*l*_3_*k*_, *l*_3_ = 1) and the number of molecules with similar temporal pattern for each molecule. Dot colors correspond to the metabolic groups (inset). Representative molecules are labeled. Their abbreviations are as follows: Cit, citrulline; CRP, C-reactive peptide; FFA, free fatty acid; GIP, gastric inhibitory polypeptide (active); Glc, glucose; Ile, isoleucine; Ins, insulin; Ketone, total ketone bodies; Leu, leucine.

In our earlier study, we defined the feature of the similarity of temporal patterns among individuals, by calculating for each molecule a correlation coefficient connecting all time courses, combining two selected from all individuals as the index of the similarity (Fig. S3, Fujita et al., 2022). The higher the value of this index of similarity, the higher the similarity of temporal patterns among individuals. Because we used *w*_*l*_3_*k*_, *l*_3_ = 1 and 2 to select the molecule with constant individual dependence (Fig. 2A), we investigated the relationship between the absolute values of both *w*_*l*_3_*k*_, *l*_3_ = 1,2 and the similarity of temporal patterns among individuals. Multiple regression analysis showed a significant relationship between the value of abs(*w*_*l*_3_*k*_, *l*_3_ = 1,2) and the index of the similarity (adjusted coefficient of determination is 0.598 (*p* = 5.75 x 10^-15^) (Fig. 4C). Thus, the value of *w*_*l*_3_*k*_, *l*_3_ = 1,2 reflects the similarity of the temporal pattern among individuals.

In our earlier study, we defined the feature of the change in the relationship among individuals over time, by calculating the average changes in z-score values over time for each molecule at each time point as the index of the change in the relationship among individuals over time (6). The larger the value of the index, the more constant the relationship among individuals over time. Because we used both *w*_*l*_3_*k*_, *l*_3_ = 2 and 4 to select molecules with variable individual dependence, we investigated the relationship between the absolute values of *w*_*l*_3_*k*_, *l*_3_ = 2,4 and the magnitude of the change in the relationship among individuals over time. Correlation analysis showed no significant relationship between the value of abs (*w*_*l*_3_*k*_, *l*_3_ = 2,4) and the magnitude of the change in the relationship among individuals over time (Table 2). We also investigated other relationships between the absolute value of *w*_*l*_3_*k*_, *l*_3_ = 1 and the magnitude of the change over time in the relationship among individuals. Correlation analysis showed a significant relationship between the value of abs(*u*_*l*_1_*i*_, *l*_1_ = 1) and the magnitude of the change in the relationship among individuals over time, but the correlation coefficient was low (*r* =3.51 x 10^-1^) (Fig. 4D, Table 2). Therefore, we conclude that TD cannot be used to extract features that reflect the magnitude of the change in the relationship among individuals over time.

In our earlier study, we defined the feature of the similarity of temporal patterns among molecules (6). Because the similarity of temporal patterns among molecules was defined for a pair of molecules, we defined the index as a quantitative value of the similarity of temporal patterns among molecules for each molecule. Molecules with large values of this index have a large number of molecules with similar temporal patterns. For details, please refer to our earlier study (6). In this study, we used *w*_*l*_3_*k*_, *l*_3_ = 1,2,4 to select molecules whose time dependence is transient or persistent. Therefore, we investigated the relationship between the absolute values of *w*_*l*_3_*k*_, *l*_3_ = 1,2,4 and the number of molecules with similar temporal patterns. Correlation analysis showed that a significant relationship between the value of abs(*w*_*l*_3_*k*_, *l*_3_ = 1) and the number of molecules with similar temporal patterns (Fig. 4E, Table 2). Therefore, the value of *w*_*l*_3_*k*_, *l*_3_ = 1 reflects the similar temporal patterns among molecules.

Taken together, the following three features were extracted by data-driven analysis: the “amplitude and rate” components of temporal patterns, the similarity of temporal patterns among individuals, and the similarity of temporal patterns among molecules. The only feature not extracted was the relationship among individuals over time, indicating that TD can extract some of the same features obtained by hypothesis-driven analysis in a non-biased manner (Figs. 4, S3, S4),

### Molecules targeted in our earlier study and extracted molecules in this study

In this study, we used TD to extract 68 molecules showing specific time dependence and individual dependence by unsupervised learning (Fig. 5, Table S4). We examined the overlap with the 83 molecules analyzed in our earlier study; the number of overlapped molecules was 55, which is 55/83 = 66% compared to the 83 molecules analyzed in our earlier study (Fig. 5, Table S4). This result indicates that our earlier hypothesis driven analysis (6) included most of the specific molecules extracted by data-driven analysis using TD in this study. Fisher’s exact test showed that the overlap was significant at a significance level of 0.01 (*p* = 3.88 x 10^-43^). The overlapped molecules included glucose and insulin, amino acids such as leucine and valine, and lipids such as FFA and total ketone body (Fig. 5, Table S4), which we analyzed in detail in our earlier study (Fujita et al., 2022). The set of molecules extracted by TD but not analyzed in our earlier study included molecules such as adrenaline, acetoacetic acid, and noradrenalin (Fig. 5, Table S4). Although these molecules may have shown specific temporal patterns, they had many missing points and were excluded from the analysis in our earlier study. Note that the TD was analyzed with missing points as zero.

**Fig. 5.**
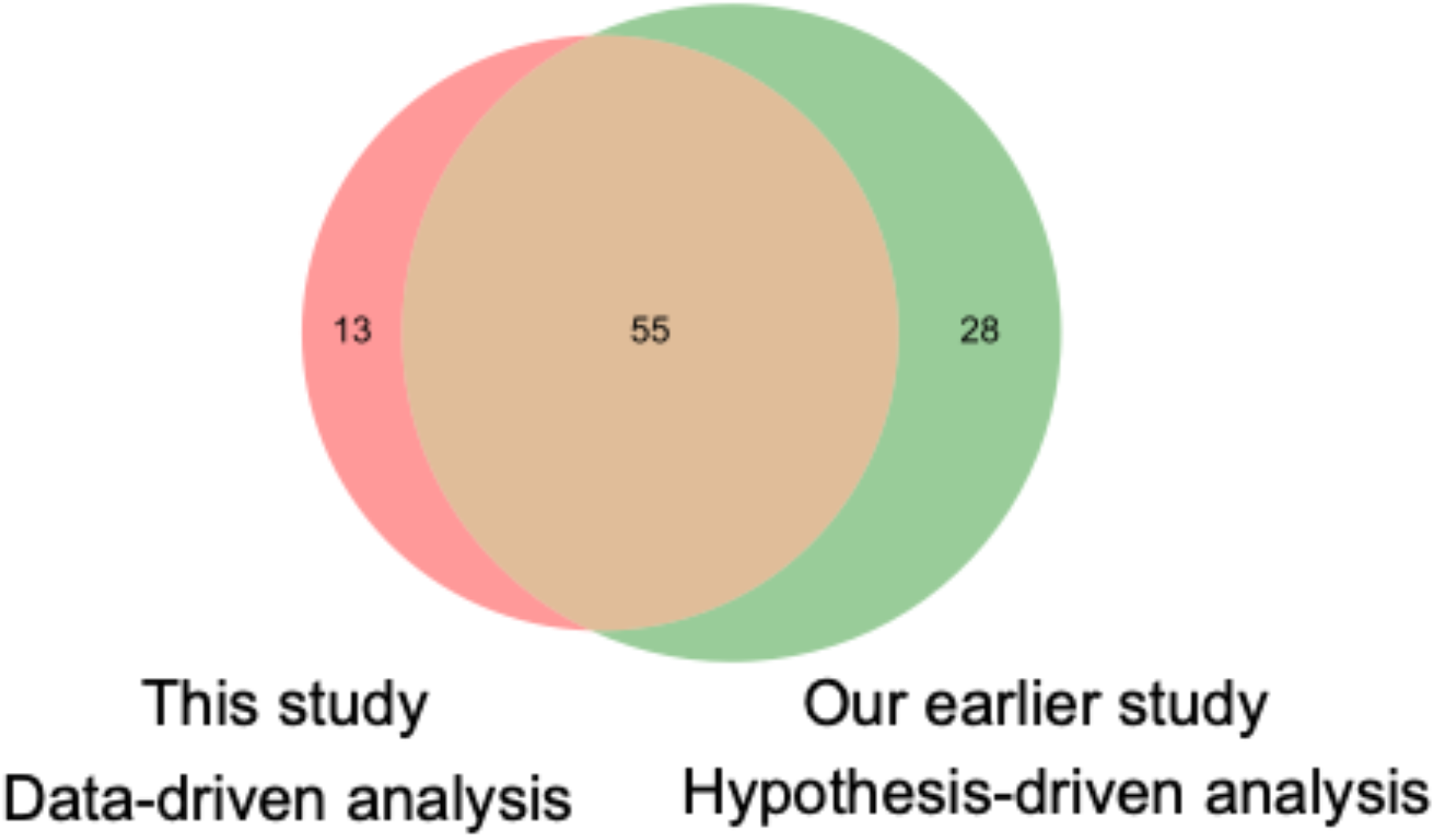
Molecules targeted in the hypothesis-driven and data-driven analyses. Venn diagram of molecules targeted in our earlier study (Fujita et al., 2022) and extracted molecules in this study. The molecules included in each Venn diagram are listed in Table S4.

By contrast, many of the molecules included in our earlier hypothesis-driven analysis but not extracted by TD were ions and other metabolites (Fig. 5, Table S4). These molecules did not show a specific temporal pattern by glucose ingestion, although they had no missing points. The overlapped 55 molecules was 55/68 = 81% compared to 68 molecules extracted using TD (Fig. 5). This result indicates that the features extracted by TD are useful in biology, as data-driven analysis sufficiently captured the features obtained by our previous hypothesis-driven analysis.

Taken together, we extracted the features of the molecules and temporal patterns that were targeted in our earlier hypothesis-driven analysis by applying TD to the datasets that were characterized in our earlier study (6). This result shows not only the validity of the results of our previous study, but also the usefulness of TD as a FE method.

### Singular vectors obtained by TD (’*fourth-order tensor with individuals mode, time mode, experimental condition mode and molecule mode*’)

We further analyzed a new dataset that additionally included differences in experimental conditions such as different doses of glucose and ingestion patterns of glucose ingestion (Table S5). For these data, three healthy volunteers ingested either three different doses of glucose (25, 50, and 75 g) rapidly or glucose over 2 h in six experiments to obtain time series data (Fig. S5) (Fujii et al., 2019). The rapid ingestion paradigm is referred to as bolus ingestion, whereas the slow ingestion paradigm is referred to as continuous ingestion over 2 h. We performed TD of four modes data of the concentrations of 40 blood molecules (4 hormones, 36 metabolites) at 26 time points in three healthy subjects by three different oral doses and two different patterns of glucose ingestion, and tried to capture the differences of temporal pattern of molecular concentration among individuals, molecules, and experimental conditions by TD.

The data were formatted as a tensor (*X* = {*x_ijkm_*} ∈ ℝ^3×26×6×40^) representing the concentration of the *i*th individual, *j*th time point, *k*th condition, and *m* th molecule. We normalized *x_ijkm_* as 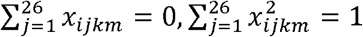. HOSVD (24) was applied to *x_ijk_*.

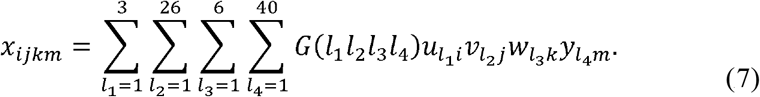

*G*(*l*_1_*l*_2_*l*_3_*l*_4_) ∈ ℝ^3×26×6×40^ is the core tensor; and *U* = {*u*_*l*_1_*i*_} ∈ ℝ^3×3^, *V* = {*v*_*l*_2_*j*_ ∈ ℝ^26×26^, *W* = *w*_*l*_3_*k*_ ∈ ℝ^6×6^, *Y* = {*y*_*l*_4_*m*_} ∈ ℝ^40×40^ are the orthogonal matrices. We sorted the core tensor components *G*(*l*_1_*l*_2_*l*_3_) in the order of their absolute values (Table 3), and focused only on the top four core tensor components and associated singular vectors (Fig. S6, Table 3). As discussed later, these four were sufficient to characterize the differences in experimental conditions. These large absolute values of *G*(*l*_1_*l*_2_*l*_3_) corresponded to the singular vectors *u*_*l*_1_*i*_, *l*_1_ = 1, *v*_*l*_2_*j*_, *l*_2_ = 1,2, *w*_*l*_3_*k*_, *l*_3_ = 1,2, *y*_*l*_4_*m*_, *l*_4_ = 1,2 of individual, time, experimental condition, and molecule (Table 3). Thus, we next interpreted these singular vectors.

*u*_*l*_1_*i*_ are the individual-related singular vectors. This value means individual dependence (Fig. 6A). *u*_*l*_1_*i*_, *l*_1_ = 1 showed a constant value among individuals (Fig. 6A). *v*_*l*_2_*j*_ are the time-related singular vectors (Fig. 6B). This value means time-dependence. *v*_*l*_2_*j*_, *l*_2_ = 1 peaked at the later time point (later than 60 min) (Fig. 6B), *v*_*l*_2_*j*_, *l*_2_ = 2 peaked at the earlier time point (at 60 min) (Fig. 6B). This result indicated that the early and late peaks were the main components of the temporal pattern.

**Fig. 6.**
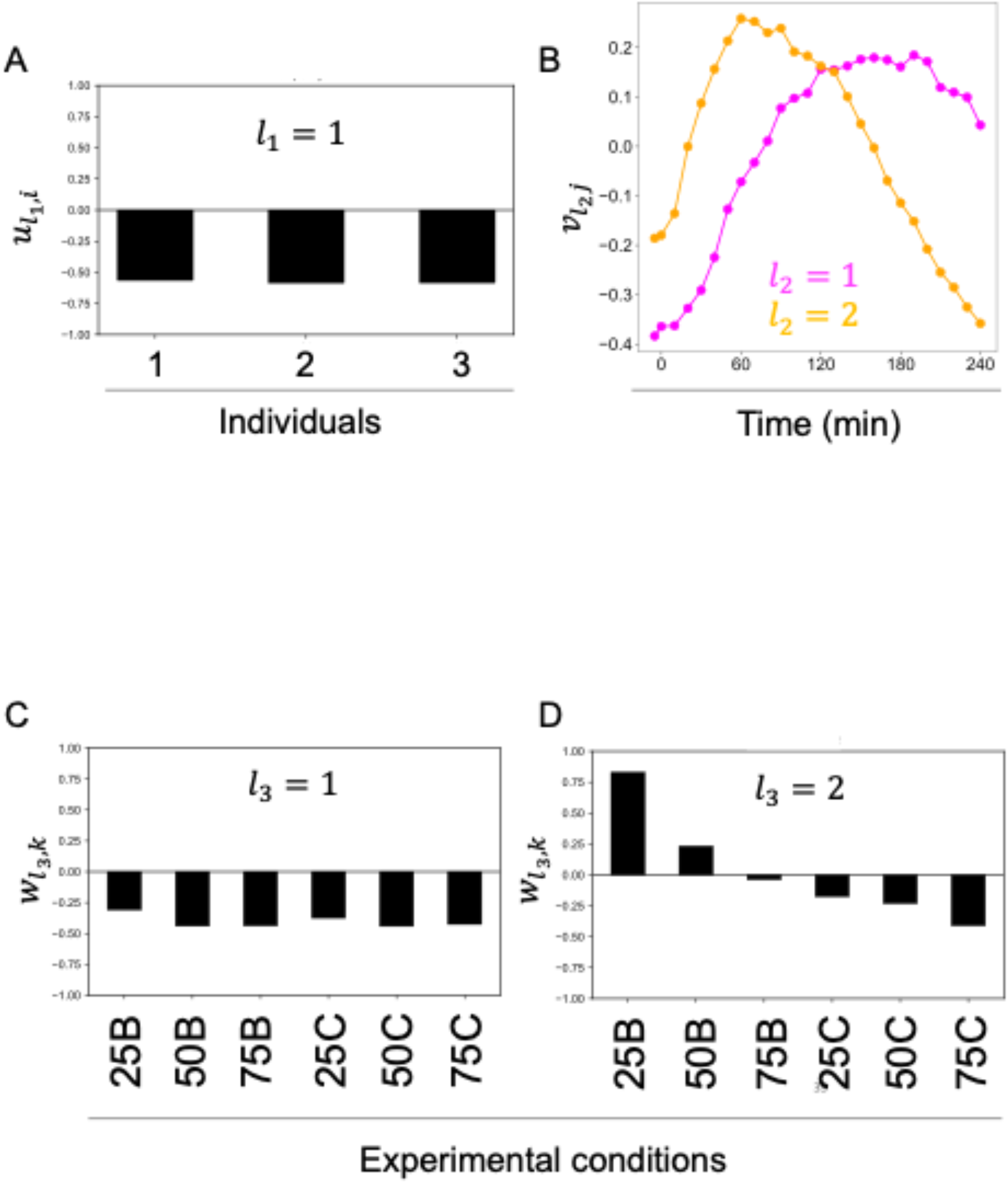
Singular vectors obtained by tensor decomposition (’*fourth-order tensor with individual mode, time mode, experimental condition mode, and molecule mode*’) **A** Individual-related singular vectors (*u*_*l*_1_*i*_, *l*_1_ = 1). **B** Time-related singular vectors (Magenta line: *v*_*l*_2_*j*_, *l*_2_ = 1 Orange liene: *v*_*l*_2_*j*_, *l*_2_ = 2). **C** Experimental condition-related singular vectors (*w*_*l*_3_*k*_, *l*_3_ = 1). D Experimental condition-related singular vectors (*w*_*l*_3_*k*_, *l*_3_ = 2). Each experimental condition is denoted by the initial letters of the amount and duration of ingestion: 25B for 25 g bolus ingestion, 75°C for 75 g-2 h continuous ingestion.

**Fig. 7.**
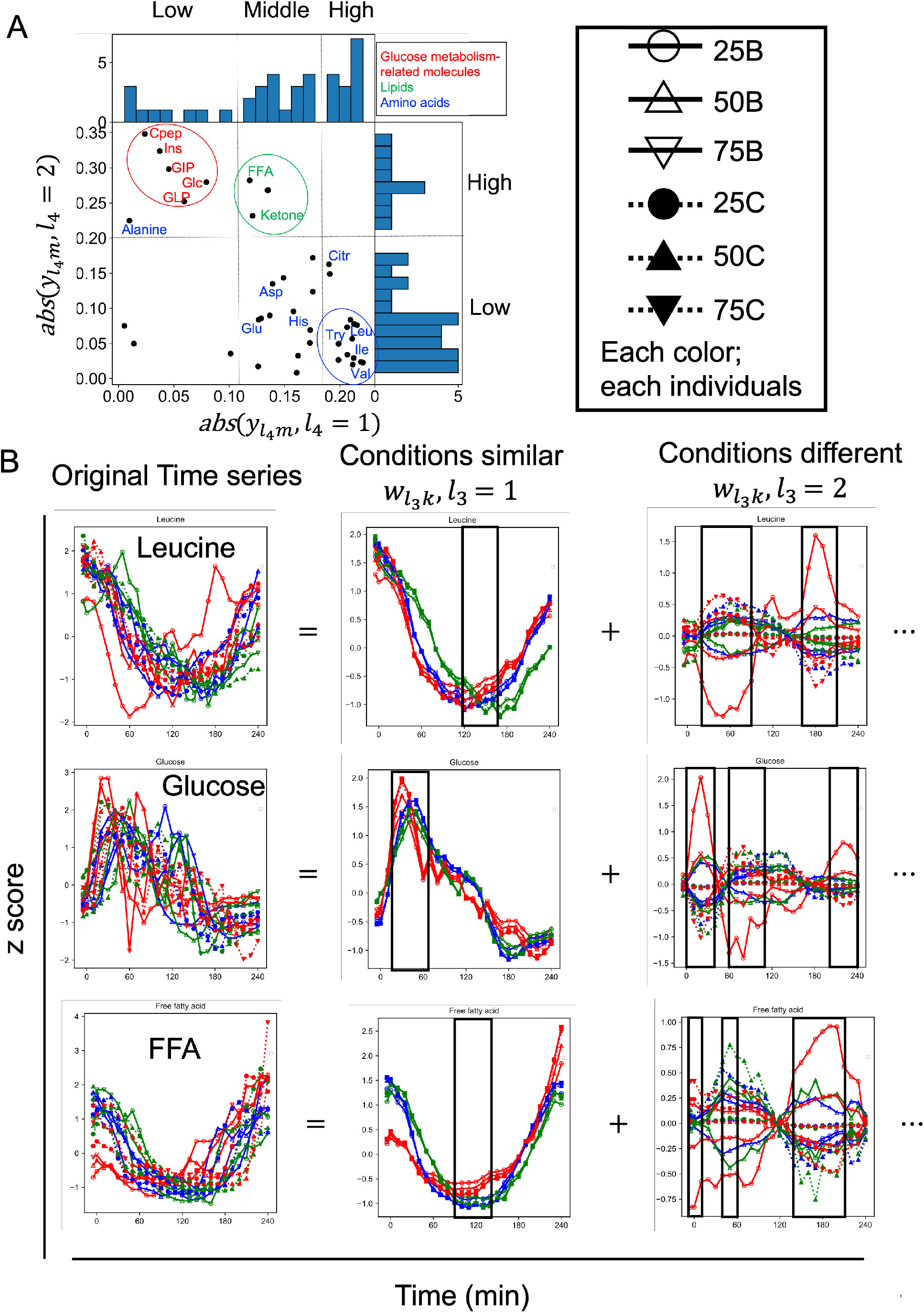
The individual-related singular vectors and decomposed time series by the dominant singular vectors (’*fourth-order tensor with individuals mode, time mode, experimental condition mode and molecule mode*’) **A** The molecule-related singular vectors (*y*_*l*_4_*m*_, *l*_4_ = 1,2). The dashed lines show the values of *y*_*l*_4_*m*_, *l*_4_ = 1 divided into three, and *y*_*l*_4_*m*_, *l*_4_ = 2 divided into two, based on the shape of the distribution. **B** Decomposed time series by the dominant singular vectors of a representative molecule and the original time series. Decomposed time series by other than the dominant singular vectors (*w*_*l*_3_*k*_, *l*_3_ > 2) are omitted. The black lines in “conditions similar” indicate the time point of the peak. The black boxes and the black lines in “conditions different” indicate the time points of high variability among conditions. The color of each line indicates each condition. Solid lines indicate bolus ingestion, and dahed lines indicate 2 h continuous ingestion. Circles, triangles, and lower triangles indicate the three doses, 25, 50, and 75 g, respectively. Abbreviations for the representative molecules are as follows: Asp aspartic acid; Cit, citrulline; CRP, C-reactive peptide; FFA, free fatty acid; 3-OH, 3-hydroxybutyric acid; Ketone, Total ketone body; GIP, gastric inhibitory polypeptide (active); Glc, glucose; GLP, glucagon-like peptide-1; Glu, glutamic acid; His, histidine; Ile, isoleucine; Ins, insulin; Leu, leucine; Tyr, tyrosine; Val, valine. The label colors correspond to the metabolic group list (inset).

*w*_*l*_3_*k*_ are the experimental condition-related singular vectors (Fig. 6C, D). This value means experimental condition dependence. *w*_*l*_3_*k*_, *l*_3_ = 1 shows the constant values among the experimental conditions (Fig. 6C). *w*_*l*_3_*k*_, *l*_3_ = 2 shows the variable values among the experimental conditions (Fig. 6D). *w*_*l*_3_*k*_, *l*_3_ = 2 also shows that the value of bolus condition decreases from positive to negative in a dose-dependent manner, and the value of continuous condition negatively increase in a dose-dependent manner. Taken together, *w*_*l*_3_*k*_, *l*_3_ = 2 divides not only the duration of the bolus or continuous condition but also the axis of the dose.

### The individual-related singular vectors and decomposed time series by the dominant singular vectors (‘*fourth-order tensor with individuals mode, time mode, experimental condition mode and molecule mode*’)

We selected the top four pairs as relatively large *G*(*l*_1_*l*_2_*l*_3_) (Fig. S6, Table 3). In previous sections (Figs. 2–5), We calculated the *p*-values for each molecule in order to select biologically meaningful molecules. However, we could not extract molecules with a significantly large value of *y*_*l*_4_*m*_ possibly because of the small number of molecules. Therefore, we summarized the features of the temporal patterns of the molecules based on the distribution of *y*_*l*_4_*m*_ (Fig. 7A).

The magnitude of the absolute value of the molecule-related singular vectors captured the features of the temporal pattern after glucose ingestion (Figs. 4, S2). We focused on the absolute value of the molecule-related singular vectors. Abs (*y*_*l*_4_*m*_) values are the absolute value of the molecule-related singular vectors. Abs (*y*_*l*_4_*m*_, *l*_4_ = 1) values of amino acids such as leucine and valine (Fig. 7A) and lipids such as FFA and ketone (Fig. 7A) were larger, and those of the molecules related to glucose metabolism such as glucose and insulin (Fig. 7A) were smaller. Abs (*y*_*l*_4_*m*_, *l*_4_ = 2) values of lipids (Fig. 7A) and glucose metabolism-related molecules (Fig. 7A) were larger, and those of amino acids were smaller (Fig. 7A). This result indicates that glucose metabolism-related molecules, amino acids and lipids have specific individual, time, and experimental condition dependence.

To better interpret the features of the temporal patterns of the molecules, we reconstructed the time series using only the singular vectors (Fig. 7B, see Methods). Because we focused on the features of the temporal patterns that were similar among experimental conditions of each molecule and the temporal patterns that were different among experimental conditions, we reconstructed the original time series of the molecule using only the constant condition dependence (*w*_*l*_3_*k*_, *l*_3_= 1) and the variable condition dependence (*w*_*l*_3_*k*_, *l*_3_ = 2) (Fig. 7B, see Methods). Here, we refer to the time series reconstructed using only (*w*_*l*_3_*k*_, *l*_3_ = 1) as “conditions similar pattern,” and the time series reconstructed using only (*w*_*l*_3_*k*_, *l*_3_ = 2) as “conditions different pattern.” The original data of leucine, FFA, and glucose, and the “conditions similar pattern” and “conditions different pattern” are shown (Fig. 7B).

Both *y*_*l*_4_*m*_, *l*_4_ = 1,2 shared a constant individual dependence (*u*_*l*_1_*i*_, *l*_1_ = 1) (Table 3). This result indicates that all molecules had a similar temporal patterns among individuals. For leucine, as one of the representative molecules of the amino acid (Fig. 7A), “conditions similar pattern” showed a later peak from 120 to 150 min (Fig. 7B). “Conditions different pattern” of leucine showed variation from 30 to 100 min and from 160 to 210 min (Fig. 7B). The earlier peak values from 30 to 100 min changed from negative to positive from bolus (Fig. 7B) to continuous (Fig. 7B), whereas later peak values from 160 to 210 min changed from negative to positive from continuous (Fig. 7B) to bolus (Fig. 7B). Because “conditions similar pattern” peaked at negative values, the more negative the value of the peak in the “Conditions difference pattern,” the larger the peak in the original time series. Thus, the earlier peak was larger for bolus, and the later peak was larger for continuous. Other amino acids with large abs (*y*_*l*_4_*m*_, *l*_4_ = 1) had constant conditional dependence (*w*_*l*_3_*k*_, *l*_3_ = 1), with time dependence of the later peak (*v*_*l*_2_*j*_, *l*_2_ = 1), whereas variable condition dependence (*w*_*l*_3_*k*_, *l*_3_ = 2) had time dependence of the earlier peak (*v*_*l*_2_*j*_, *l*_2_ = 2) (Table 3). This result suggested that the temporal patterns of amino acids showed similar temporal patterns for the later peak and different temporal patterns for the earlier peak among the experimental conditions.

For glucose, the representative molecule of glucose metabolism-related molecules, “conditions similar pattern” showed an early peak from 30 to 40 min (Fig. 7B). “Conditions different pattern” showed variation from 0 to 40 min, 60 to 110 min, and 200 to 240 min, but the variation from 200 to 240 min was smaller than the other two durations (Fig. 7B). During 0 to 40 min and 200 to 240 min, the peak value changed from negative to positive from continuous (Fig. 7B) to bolus (Fig. 7B), whereas from 60 to 110 min, the peak value changed from negative to positive from bolus (Fig. 7B) to continuous (Fig. 7B). Because “conditions similar pattern” peaked at positive values, this result indicates that the earlier peak during 0 to 40 min was larger for bolus and the later peak during 60 to 110 min was larger for continuous. This result also indicates that the other peak during 200 to 240 min after glucose ingestion was larger for bolus although the variation among experimental conditions was not as large as the others. Glucose metabolism-related molecules such as glucose and insulin with large abs (*y*_*l*_4_*m*_, *l*_4_ = 2) had constant experimental condition dependence (*w*_*l*_3_*k*_, *l*_3_ = 1) with time dependence of the early peak (*v*_*l*_2_*j*_, *l*_2_ = 2) (Table 3), but variable experimental condition dependence (*w*_*l*_3_*k*_, *l*_3_ = 2) with the time dependence of the late peak (*v*_*l*_2_*j*_, *l*_2_ = 1) (Table 3). These results suggest that the temporal patterns of glucose metabolism-related molecules showed similar temporal patterns for the earlier peak and different temporal patterns for the later peak among the experimental conditions.

For FFA, as one of the representative molecules of the lipid, “conditions similar pattern” peaked from 110 to 120 min, later than those of glucose metabolism-related molecules but earlier than amino acids (Fig. 7A). “Conditions different pattern” showed variation from −5 to 10 min, 40 to 60 min, and 140 to 210 min (Fig. 7B). The peak values changed from negative to positive from bolus (Fig 7B) to continuous (Fig. 7B) from 5 to 10 min, 40 to 60 min, and from negative to positive from continuous to bolus during 140 to 210 min. Because “conditions similar pattern” peaked at negative values, this result indicates that the earlier peak was larger for bolus and the later peak was larger for continuous. Abs (*y*_*l*_4_*m*_, *l*_4_ = 1) of lipids were also larger than those of glucose metabolism-related molecules but smaller than those of amino acids (Fig. 7A). In addition, abs (*y*_*l*_4_*m*_, *l*_4_ = 2) of lipids were large (Fig. 7A). This result suggests that the similar temporal patterns among experimental conditions for lipids showed a later peak than glucose metabolism-related molecules and earlier peak than amino acids. This result also suggests that the different temporal patterns among experimental conditions for lipids showed an earlier peak than glucose metabolism-related molecules and later peak than amino acids.

Taken together, the similar temporal patterns among experimental conditions changed from the earlier peak to later peak in the order of amino acids, lipids, and glucose metabolism-related molecules (Fig. 7, Table 3). “Conditions similar patterns” of leucine, FFA, and glucose showed these time dependencies (Fig. 7B). The molecule-related singular vectors of each molecule also indicated that the different temporal patterns among experimental conditions changed from the earlier peak to the late peak in the order of amino acids, lipids, and glucose metabolism-related molecules (Fig. 7, Table 3). “Conditions different pattern” for leucine, FFA, and glucose showed large variation at different time points (Fig. 7B). “Conditions different pattern” also showed that the earlier peak was larger for bolus and the later peak was larger for continuous (Fig. 7B). This supports that the change in *w*_*l*_3_*k*_, *l*_3_ = 2 from positive to negative from 25B to 75C indicated a change from the earlier peak to the later peak. Taken together, the temporal patterns of each molecule were characterized by experimental condition dependence (constant or variable) and time dependence (early or late peak).

## DISCUSSION

### The temporal pattern of *‘third-order tensor with individual mode, time mode, and molecule mode’*

In this study, we applied TD to *‘third-order tensor with individual mode, time mode, and molecule mode’* as multimodal data, and extracted features of temporal patterns obtained by previous data-driven analysis (Figs. 1–5, Tables 1, 2). In our earlier study (6), we selected target molecules from the same dataset and characterized four features of temporal patterns of molecules among individuals and molecules derived from the hypothesis-driven analysis.

The features extracted by the TD in this study reflected three of the four features of the temporal pattern (Figs. 4, S3, S4). This result suggests that not only the three features of the temporal pattern capture the features of the dataset but also that a feature can only be extracted by hypothesis-driven analysis. However, we could not exclude the possibility that the lower core tensor components reflect such feature, and further study is necessary to address this issue.

We also extracted 68 molecules that showed specific time and individual dependencies through unsupervised learning by using TD (Fig. 5). The extracted molecules included most of the molecules analyzed in our earlier study, which ensured the validity of the extraction of feature by our earlier hypothesis-driven study (Fujita et al., 2022). In addition, the features extracted by TD sufficiently captured the features of the temporal patterns of the molecules that were of interest in our previous study, indicating that the results of data-driven analysis are useful in biology.

### Interpretation of time patterns by reconstructing time series

To better interpret the features of the temporal patterns of molecules, we reconstructed the time series using only the singular vectors of interest (Figs. 3,7, see Methods). For example, a molecule such as glucose, which has multiple time and individual dependencies (Fig. 3), is difficult to extract multiple biological features. We improved the interpretability by reconstructing the time series and visualizing it (Fig. 3B). In the field of biology, the extraction of waveforms by low-rank approximation in functional magnetic resonance imaging has been studied (30). In our earlier study, the low-rank representation of the time series facilitated biological interpretation and led to useful discussions (Fujita et al., 2022).

### The temporal pattern of ‘*fourth-order tensor with individual mode, time mode, experimental condition mode, and molecule mode*’

We also applied TD to ‘*fourth-order tensor with individual mode, time mode, experimental condition mode, and molecule mode*’ to characterize the experimental condition-dependent temporal patterns for each molecule (Fig. 6). The similar and different temporal patterns among experimental conditions changed from the earlier peak to the later peak in the order of amino acids, lipids, and glucose metabolism-related molecules (Fig. 7, Table 3).

“Conditions different pattern” of leucine, glucose, and FFA showed a large variation at different time points. “Conditions different pattern” of leucine, glucose, and FFA showed that the earlier peaks were larger for bolus ingestion and the later peaks were larger for continuous ingestion. In a previous study in which healthy subjects drank the same dose of glucose solution in either a bolus or continuous manner, the peaks by continuous ingestion were later than those by bolus ingestion for the glucose metabolism-related molecules such as glucose, insulin, C-peptide, and GIP (31); branched-chain amino acids such as valine, leucine, and isoleucine; and metabolites including FFAs and hormones (32). The results of this study were consistent with previous studies in which the dependence of the experimental conditions was evaluated for each molecule (31,32). However, in this study, we could evaluate not only the dependence among experimental conditions for each molecule (Fig. 7B) but also the difference in dependence of the experimental conditions among molecules (Fig. 7A). This indicates that the analysis flow using TD is valid for multimodal data and can simultaneously extract multiple biological features by data-driven analysis.

### Research limitations and prospects

We focused on the top four pairs as the core tensors with the largest absolute values (Tables 1, 3). However, because this selection method introduces analyst bias, selection criteria will be needed in the future. In addition, a few molecules possibly had a specific dependence on the core tensor that we did not focus on in this study. A decomposition method that can extract the feature dependence of a few molecules will be necessary. The various features of the responses of molecules (temporal patterns) by glucose ingestion we focused on in this study were clarified by a combination of hypothesis-driven and data-driven analyses (Fujita et al., 2022). However, we could not identify the mechanisms behind the feature of dataset. For this purpose, it would be effective to formulate hypotheses based on the features revealed in this study and to make plans for new experiments or analyze time series data using mechanistic mathematical models. We discussed interindividual differences in the role of incretins in the regulation of blood glucose levels by mathematical model analysis focusing on glucose metabolism-related molecules (26).

### Conclusion

In this study, we first applied TD to a dataset representing changes in the concentrations of 562 molecules (7 hormones and 555 metabolites) at 14 time points in 20 healthy subjects by 75 g glucose ingestion (‘*third-order tensor with individual mode, time mode, and molecule mode*’). We obtained the core tensor and individual-, time-, and molecule-related singular vectors. By reconstructing the time series using only the singular vectors that we focused on, we could better interpret the features of the temporal patterns of the molecules. We characterized each molecule by individual dependence (constant or variable) and time dependence (sustained or transient).

The molecule-related singular vectors obtained by TD reflected three of the four features characterized by our earlier hypothesis-driven study (Fujita et al., 2022). We also extracted 68 molecules showing feature time and individual dependencies through unsupervised learning method by using TD. The extracted molecules overlapped significantly with the analyzed molecules in our earlier study (6). Therefore, by applying TD to the dataset characterized in our earlier study, we extracted the features of the target molecules and the revealed temporal patterns in our earlier study (6). This result not only confirms the validity of the results of our earlier study (6) but also shows the usefulness of TD as a FE method.

Next, we applied the TD method to a dataset representing the concentration changes of 40 molecules in three healthy subjects at 26 time points by three different oral doses and two different patterns of glucose ingestion (‘*fourth-order tensor with individual mode, time mode, experimental condition mode, and molecule mode*’). We obtained the core tensor and individual-, time-, molecule-, and experimental condition-related singular vectors. By reconstructing the time series using only the singular vectors we focused on, we could better interpret the features of the temporal patterns of the molecules. We can characterize the temporal pattern of each molecule by its experimental condition dependence (constant or variable) and time dependence (early or late peaks). We applied TD to a time series dataset of human blood metabolites and hormone concentrations before and after glucose ingestion with various modes, and extracted biological features in a non-biased manner without time-consuming process of hypothesis generation. We propose that TD can be the first choice for analysis of omics data.

## CONTACT FOR REAGENT AND RESOURCE SHARING

Further information and requests for resources and reagents should be directed to and will be fulfilled by the Lead Contact, Shinya Kuroda.

## SUPPLEMENTAL INFORMATION

Supplemental Information includes 6 figures and 5 tables.

## Data and Code Availability

All data generated or analysed for *‘third-order tensor with individual mode, time mode, and molecule mode’* is included in the article (6) and its supplementary materials files. The data for *‘fourth-order tensor with individual mode, time mode, experimental condition mode, and molecule mode’* is included in this article and its supplementary materials files. The code files used in the simulation are freely available at https://github.com/sfujita0601/TDA_HumanOGTT

## ACKNOWLEDGEMENTS

We thank our laboratory members for critical reading of the manuscript.

## AUTHOR CONTRIBUTIONS

S.F., K.H., Y.T. and S.K. analysed the data. S.F., Y.K., and S.K. wrote the manuscript. The study was conceived and supervised by S.F. and S.K.

## REFERENCES

1. Nielsen J. Systems biology of metabolism. Annu Rev Biochem. 2017;86:245–75.

2. Shi-kai YAN, Run-hui LIU, Hui-zi JIN, Xin-ru LIU, Ji YE, Lei S. “Omics” in pharmaceutical research□: overview, applications, challenges, and future perspectives. 2015;13(2011):3–21.

3. Conesa A, Prats-montalbán JM, Tarazona S, José M, Ferrer A. Chemometrics and Intelligent Laboratory Systems A multiway approach to data integration in systems biology based on Tucker3 and N-PLS. Chemom Intell Lab Syst [Internet]. 2010;104(1):101–11. Available from: http://dx.doi.org/10.1016/j.chemolab.2010.06.004

4. Andrikopoulos S, Blair AR, Deluca N, Fam BC, Proietto J. Evaluating the glucose tolerance test in mice. Am J Physiol Metab [Internet]. 2008 Dec;295(6):E1323–32. Available from: https://www.physiology.org/doi/10.1152/ajpendo.90617.2008

5. Bar-Joseph Z, Gitter A, Simon I. Studying and modelling dynamic biological processes using time-series gene expression data. Nat Rev Genet. 2012;13(8):552–64.

6. Fujita S, Karasawa Y, Fujii M, Hironaka K, Uda S, Kubota H, et al. Four features of temporal patterns characterize similarity among individuals and molecules by glucose ingestion in humans. npj Syst Biol Appl [Internet]. 2022 Dec 8;8(1):6. Available from: https://www.nature.com/articles/s41540-022-00213-0

7. Taguchi YH. One-class Differential Expression Analysis using Tensor Decomposition-based Unsupervised Feature Extraction Applied to Integrated Analysis of Multiple Omics Data from 26 Lung Adenocarcinoma Cell Lines. Proc - 2017 IEEE 17th Int Conf Bioinforma Bioeng BIBE 2017. 2017;2018-Janua:131–8.

8. Shaham O, Wei R, Wang TJ, Ricciardi C, Lewis GD, Vasan RS, et al. Metabolic profiling of the human response to a glucose challenge reveals distinct axes of insulin sensitivity. Mol Syst Biol [Internet]. 2008 Aug 5;4(214):1–9. Available from: http://msb.embopress.org/cgi/doi/10.1038/msb.2008.50

9. Sun WW, Hao B, Li L. Tensors in Modern Statistical Learning Background. 2019;(2009):1–34.

10. Hériché J-K, Alexander S, Ellenberg J. Integrating Imaging and Omics: Computational Methods and Challenges. Annu Rev Biomed Data Sci. 2019;2(1):175–97.

11. Taguchi YH. Identification of candidate drugs using tensor-decomposition-based unsupervised feature extraction in integrated analysis of gene expression between diseases and DrugMatrix datasets. Sci Rep. 2017;7(1):1–13.

12. Taguchi Y, Turki T. Novel method for the prediction of drug-drug Interaction based on gene expression profiles. Eur J Pharm Sci [Internet]. 2021;160(January):105742. Available from: https://doi.org/10.1016/j.ejps.2021.105742

13. Taguchi YH, Turki T. Tensor Decomposition-Based Unsupervised Feature Extraction Applied to Single-Cell Gene Expression Analysis. Front Genet. 2019;10(September):1–11.

14. Taguchi YH. Tensor decomposition-based unsupervised feature extraction applied to matrix products for multi-view data processing. Vol. 12, PLoS ONE. 2017. 1–36 p.

15. Taguchi YH, Turki T. Unsupervised tensor decomposition-based method to extract candidate transcription factors as histone modification bookmarks in post-mitotic transcriptional reactivation. Chernov A, editor. PLoS One [Internet]. 2021 May 25;16(5 May):e0251032. Available from: https://dx.plos.org/10.1371/journal.pone.0251032

16. Taguchi Y, Turki T. Tensor-Decomposition-Based Unsupervised Feature Extraction in Single-Cell Multiomics Data Analysis. Genes (Basel) [Internet]. 2021 Sep 18;12(9):1442. Available from: https://www.mdpi.com/2073-4425/12/9/1442

17. Acar E, Bro R, Smilde AK. Data Fusion in Metabolomics Using Coupled Matrix and Tensor Factorizations. Proc IEEE [Internet]. 2015 Sep;103(9):1602–20. Available from: http://ieeexplore.ieee.org/document/7202834/

18. Alter O, Golub GH. Reconstructing the pathways of a cellular system from genome-scale signals by using matrix and tensor computations. 2005;102(49).

19. Dyrby M, Baunsgaard D, Bro R, Engelsen SB. Multiway chemometric analysis of the metabolic response to toxins monitored by NMR. Chemom Intell Lab Syst. 2005;76(1):79–89.

20. Gardlo A, Smilde AK, Hron K, Hrdá M, Karlíková R, Friedecký D, et al. Normalization techniques for PARAFAC modeling of urine metabolomic data. Metabolomics. 2016;12(7).

21. Yahyanejad F. Higher order analysis of gene correlations by tensor decomposition. bioRxiv. 2019;1–22.

22. Fanaee-T H, Thoresen M. Multi-insight visualization of multi-omics data via ensemble dimension reduction and tensor factorization. Hancock J, editor. Bioinformatics [Internet]. 2019 May 15;35(10):1625–33. Available from: https://academic.oup.com/bioinformatics/article/35/10/1625/5116143

23. Yener B, Acar E, Aguis P, Bennett K, Vandenberg SL, Plopper GE. Multiway modeling and analysis in stem cell systems biology. BMC Syst Biol [Internet]. 2008 Dec 14;2(1):63. Available from: https://bmcsystbiol.biomedcentral.com/articles/10.1186/1752-0509-2-63

24. Taguchi Y. Unsupervised Feature Extraction Applied to Bioinformatics [Internet]. Cham: Springer International Publishing; 2020. (Unsupervised and Semi-Supervised Learning). Available from: http://link.springer.com/10.1007/978-3-030-22456-1

25. Sano T, Kawata K, Ohno S, Yugi K, Kakuda H, Kubota H, et al. Selective control of up-regulated and down-regulated genes by temporal patterns and doses of insulin. Sci Signal. 2016;9(455):1–12.

26. Fujii M, Murakami Y, Karasawa Y, Sumitomo Y, Fujita S, Koyama M, et al. Logical design of oral glucose ingestion pattern minimizing blood glucose in humans. npj Syst Biol Appl [Internet]. 2019 Dec 2;5(1):31. Available from: http://dx.doi.org/10.1038/s41540-019-0108-1

27. Ishii N, Nakahigashi K, Baba T, Robert M, Soga T, Kanai A, et al. Multiple high-throughput analyses monitor the response of E. coli to perturbations. Science (80-) [Internet]. 2007 Apr 27;316(5824):593–7. Available from: https://www.science.org/doi/10.1126/science.1132067

28. Soga T, Baran R, Suematsu M, Ueno Y, Ikeda S, Sakurakawa T, et al. Differential Metabolomics Reveals Ophthalmic Acid as an Oxidative Stress Biomarker Indicating Hepatic Glutathione Consumption. J Biol Chem [Internet]. 2006 Jun 16;281(24):16768–76. Available from: https://pubs.acs.org/sharingguidelines

29. Soga T, Igarashi K, Ito C, Mizobuchi K, Zimmermann HP, Tomita M. Metabolomic profiling of anionic metabolites by capillary electrophoresis mass spectrometry. Anal Chem. 2009;81(15):6165–74.

30. Hunyadi B, Dupont P, Van Paesschen W, Van Huffel S. Tensor decompositions and data fusion in epileptic electroencephalography and functional magnetic resonance imaging data. Wiley Interdiscip Rev Data Min Knowl Discov. 2017;7(1):1–15.

31. Jenkins DJA, Wolever TMS, Ocana AM, Vuksan V, Cunnane SC, Jenkins M, et al. Metabolic Effects of Reducing Rate of Glucose Ingestion by Single Bolus Versus Continuous Sipping. Diabetes [Internet]. 1990 Jul 1;39(7):775–81. Available from: http://diabetes.diabetesjournals.org/cgi/doi/10.2337/diab.39.7.775

32. Heine RJ, Hanning I, Morgan L, Alberti KGMM. The Oral Glucose Tolerance Test (OGTT): Effect of Rate of Ingestion of Carbohydrate and Different Carbohydrate Preparations. Diabetes Care [Internet]. 1983 Sep 1;6(5):441–5. Available from: http://care.diabetesjournals.org/cgi/doi/10.2337/diacare.6.5.441

