## Supplementary material for "Features extracted using tensor decomposition reflect the biological features of the temporal patterns of human blood multimodal metabolome": N.A.

### Supplementary Figures

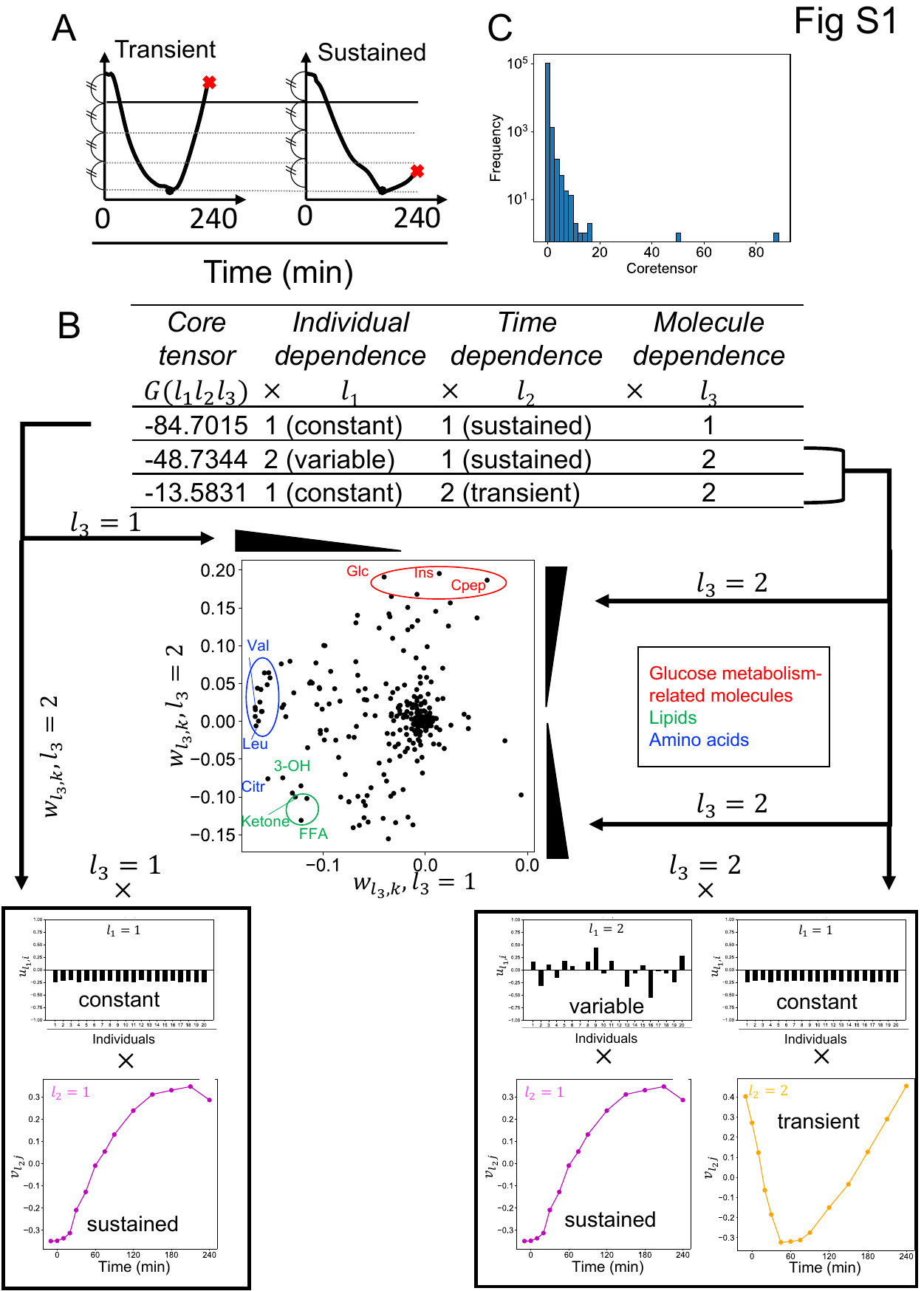

Fig. S1 Procedures for selecting a set of molecules with specific time- or individual-dependent properties

**A** Definition of transient and sustained temporal patterns. **B** Procedures for selecting a set of molecules with specific time-dependent or individual-dependent properties **C** Distribution of absolute values of the core tensor.

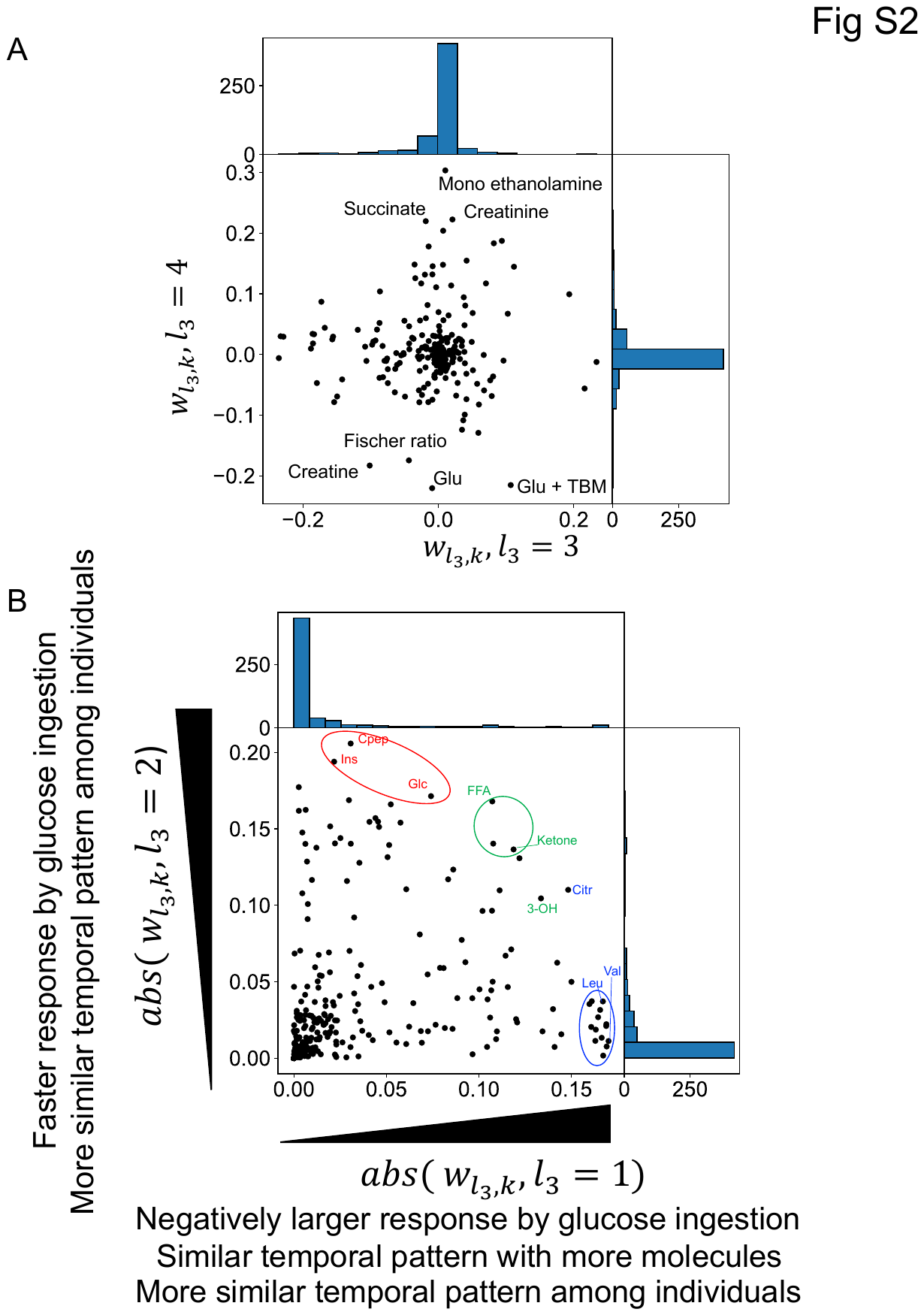

#### Fig. S2 The absolute value of the molecule-related singular vectors

**A** The distribution of the molecule-related singular vectors ($w_{l_{3}k}{,l}_{3}=1,$2). **B** The distribution of the absolute value of the molecule-related singular vectors ($w_{l_{3}k}{,l}_{3}=1,$2). Representative molecules are labeled. Abbreviations for the representative molecules are as follows: Cit, citrulline; CRP, C-reactive peptide; FFA, free fatty acid; 3-OH, 3-hydroxybutyric acid; Ketone, Total ketone body; Glc, glucose; Ins, Glu, glutamic acid; insulin; Leu, leucine; Val, valine. The label colours correspond to the metabolic group list.

#### Fig. S3 Four features of the temporal pattern of molecules

In our earlier study, we used hypothesis-driven analysis and characterized the temporal patterns among individuals and among molecules by the hypothesis-driven analysis with four features (Fujita et al., 2022): the decomposability into “amplitude” and “rate” components, the similarity of temporal patterns among individuals, the relationship among individuals’ over time, and the similarity of temporal patterns among molecules.

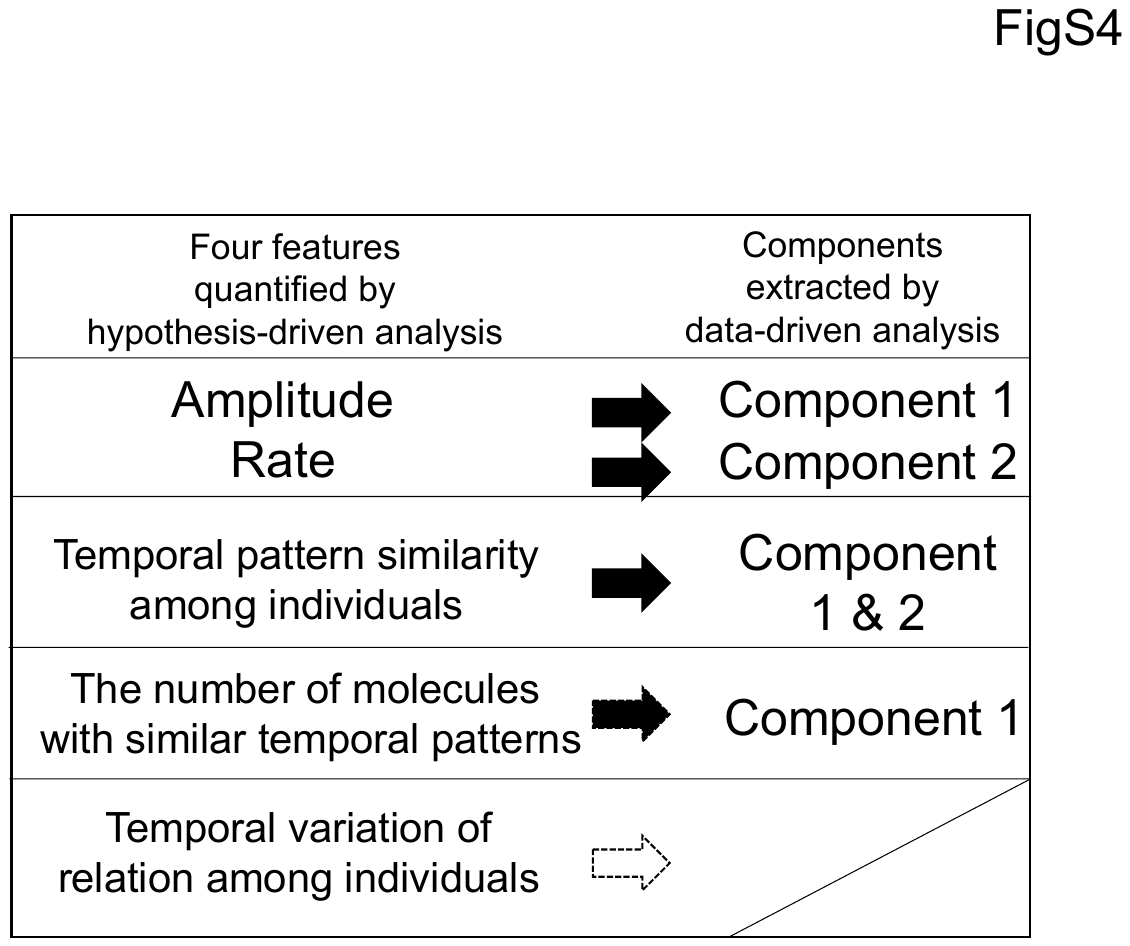

#### Fig. S4 Relationships between the four features derived from the hypothesis-driven analyais and molecule components extracted from data-driven analysis

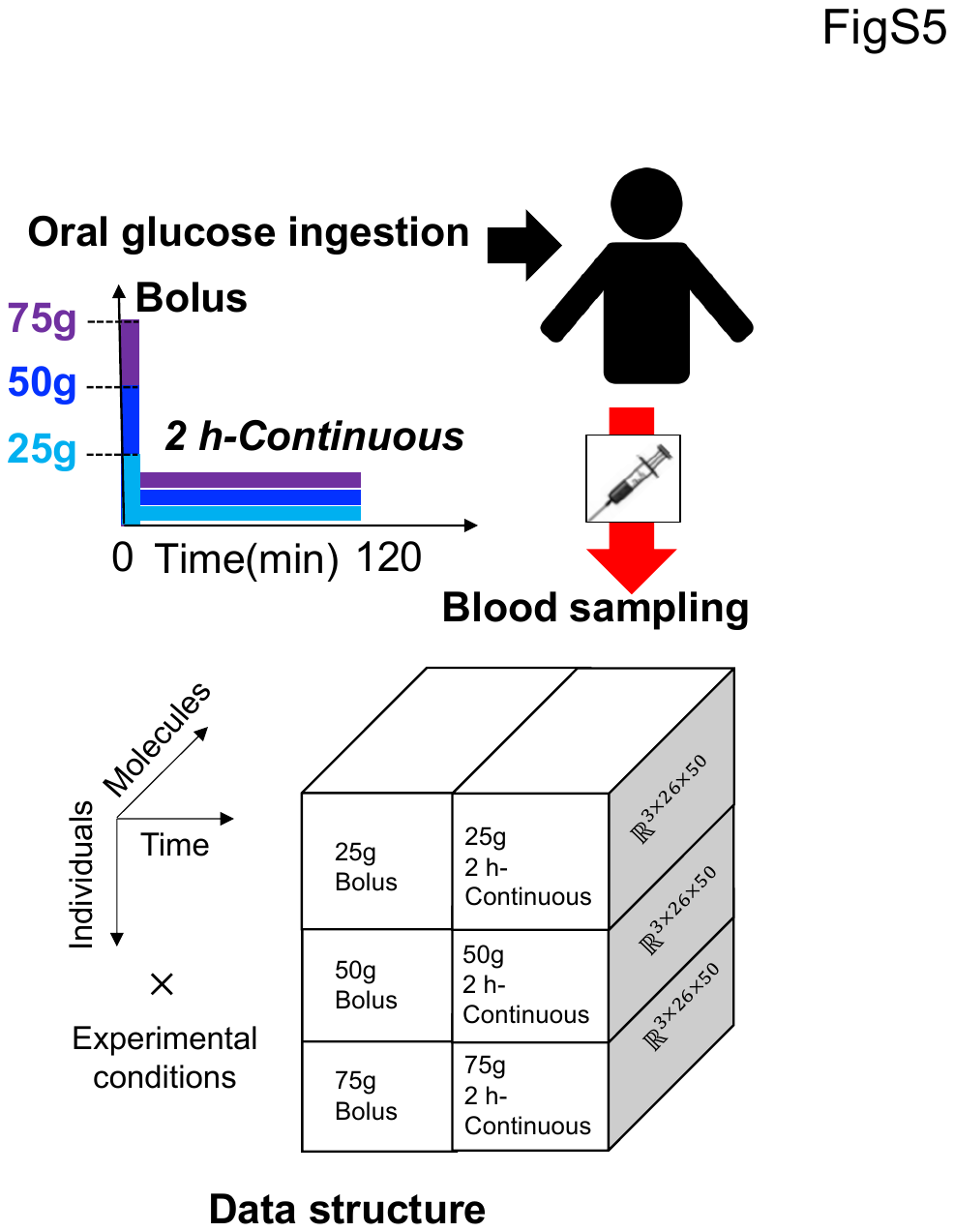

#### Fig. S5 ‘fourth-order tensor with individuals mode, time mode, experimental condition mode and molecule mode’

Three subjects orally ingested glucose with three doses 75, 50, and 25 g in two durations of bolus and 2 h continuous ingestion. The data structure has four axes: individual $\times$ time $\times$ experimental condition $\times$molecule. The data represent the concentration changes at 26 time points (-5, 0, 10, 20, 30, 40, 50, 60, 70, 80, 90, 100, 110, 120, 130, 140, 150, 160, 170, 180, 190, 200, 210, 220, 230, 240 minutes) from 5 min before fasting to 240 min after glucose ingestion for 40 molecules in three healthy subjects, in six different experimental conditions.

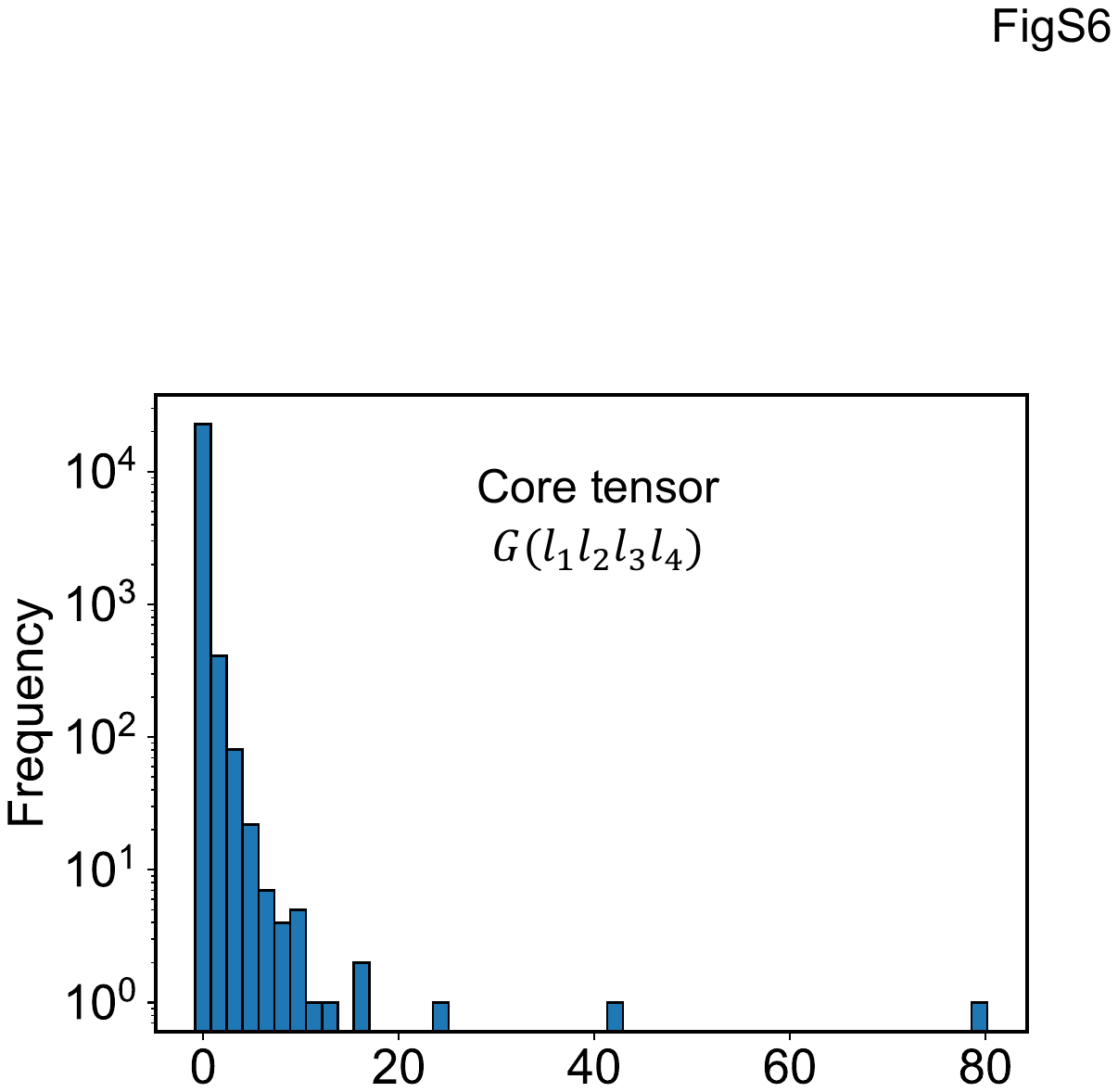

#### Fig. S6 The distribution of absolute values of the core tensor（’***fourth-order tensor with individuals mode, time mode, experimental condition mode and molecule mode***’）
